# Conserved and novel enhancers in the *Aedes aegypti single-minded* locus recapitulate embryonic ventral midline gene expression

**DOI:** 10.1101/2023.08.01.551414

**Authors:** Isabella Schember, William Reid, Geyenna Sterling-Lentsch, Marc S. Halfon

## Abstract

Transcriptional *cis*-regulatory modules, e.g., enhancers, control the time and location of metazoan gene expression. While changes in enhancers can provide a powerful force for evolution, there is also significant deep conservation of enhancers for developmentally important genes, with function and sequence characteristics maintained over hundreds of millions of years of divergence. Not well understood, however, is how the overall regulatory composition of a locus evolves, with important outstanding questions such as how many enhancers are conserved vs. novel, and to what extent are the locations of conserved enhancers within a locus maintained? We begin here to address these questions with a comparison of the respective *single-minded (sim)* loci in the two dipteran species *Drosophila melanogaster* (fruit fly) and *Aedes aegypti* (mosquito). *sim* encodes a highly conserved transcription factor that mediates development of the arthropod embryonic ventral midline. We identify two enhancers in the *A. aegypti sim* locus and demonstrate that they function equivalently in both transgenic flies and transgenic mosquitoes. One *A. aegypti* enhancer is highly similar to known *Drosophila* counterparts in its activity, location, and autoregulatory capability. The other differs from any known *Drosophila sim* enhancers with a novel location, failure to autoregulate, and regulation of expression in a unique subset of midline cells. Our results suggest that the conserved pattern of *sim* expression in the two species is the result of both conserved and novel regulatory sequences. Further examination of this locus will help to illuminate how the overall regulatory landscape of a conserved developmental gene evolves.

**AUTHOR SUMMARY:** The expression patterns and roles of genes, especially those involved in core developmental processes, are often conserved over vast evolutionary distances. Paradoxically, the DNA sequences surrounding these genes, which contain the *cis*-regulatory sequences (enhancers) that regulate gene expression, tend to be highly diverged. The manner and extent to which enhancers are functionally conserved, and how the overall organization of regulatory sequences within a locus is preserved or restructured, is not well understood. In this paper, we investigate these questions by identifying enhancers controlling expression of a master nervous system regulatory gene named *sim* in the mosquito *Aedes aegypti*, and comparing their functions and locations to those in the well-characterized *sim* locus of the fruit fly *Drosophila melanogaster*. Our results suggest that the two species generate identical patterns of *sim* expression through a mix of conserved and novel regulatory sequences. Continued exploration of the *sim* locus in these two species will help to build a comprehensive picture of how a regulatory locus for a master developmental regulator has evolved.

## INTRODUCTION

Transcriptional *cis*-regulatory modules (CRMs)—e.g., enhancers and silencers—are essential functional elements necessary for maintaining proper gene expression throughout all stages of the life cycle (1). They also serve as drivers of evolutionary change, enabling gains and losses of gene activity in specific cells and tissues and reconfiguring the gene regulatory networks (GRNs) governing development and differentiation (2). However, it is also apparent that conserved GRNs and developmental processes can utilize functionally conserved CRMs, which retain functional and mechanistic homology over hundreds of millions of years of evolution (e.g. 3, 4-11). In these cases, the DNA sequence of the CRMs has often diverged well past the point of possible linear alignment, but non-linear alignment methods or (in some cases) analysis of common transcription factor binding sites allow for their identification. Still poorly understood, however, is how the overall regulatory makeup of a locus changes over evolutionary time. In cases where this has been studied, some functionally homologous enhancers seem to be maintained in orthologous positions, whereas others appear to be “nomadic” and ply their functions from new genomic locations (4, 12, 13). Not known is how many of the CRMs contributing to a gene’s overall expression pattern are conserved, regardless of location, or what the potential ratio of conserved:novel enhancers tends to be. Existing analyses are complicated by a number of factors, including the frequent presence of semi-redundant “shadow” enhancers (14), which makes it difficult to determine if the CRMs being compared are lineally related; different evolutionary timescales with varying degrees of genomic sequence divergence; functionally distinct types of gene loci, with some genes mediating essential core regulatory functions (e.g., *Drosophila twist*, a main regulator of gastrulation (13)) whereas others are involved in more differentiated traits showing significant phenotypic variation (e.g., *Drosophila yellow*, involved in pigmentation (12, 15)); and lack of reciprocal testing of putative enhancers, making it difficult to be certain that they are functionally equivalent in their respective native genomes.

We have been studying the evolution of the insect *single-minded (sim)* locus to gain insight into how the regulatory landscape of a conserved developmental gene locus evolves. *sim,* which encodes a bHLH-PAS family transcription factor, is the primary regulatory gene mediating development of the midline of the arthropod ventral nervous system, with conserved early midline expression observed throughout the Arthropoda from flies to spiders (16–21). Like its vertebrate counterpart the floor plate, the midline of the arthropod nerve cord is a specialized structure critical for normal development and an important source of inductive signals and axon guidance molecules (22). *sim* has been studied extensively in *Drosophila melanogaster* (FlyBaseID: FBgn0004666), where it initiates expression prior to gastrulation in the mesectoderm, two single-cell wide rows abutting the embryonic presumptive mesoderm (“mesectoderm” stage). *sim* expression persists in all midline cells through germband retraction (“midline primordium” stage), as well as in a small subset of somatic muscles. Following germband retraction, *sim* is maintained in a subset of the midline lineage (“late midline” stage) (16, 23, 24). Sim binding sites have been identified in almost all known midline CRMs, and Sim binding contributes to midline gene expression in all cases tested to date (25–34). In post-embryonic stages of development, *sim* expression is observed in the brain and ovaries (35, 36).

Regulation of *Drosophila sim* has been explored in detail, with virtually all of the non-coding DNA in the *sim* locus tested for regulatory activity in reporter gene assays (31, 35, 37–40). At least four midline primordium stage enhancers have been identified, one upstream of the gene and three within the large first intron of the *sim-RB/RD* transcripts. CRMs for mesectoderm-stage *sim* expression and for larval and adult *sim* expression have been defined as well (35, 37). Here, we describe the identification of two midline primordium enhancers in the *sim* locus of the mosquito *Aedes aegypti*. Our choice of *A. aegypti* was originally motivated by previous reports that *sim* expression undergoes a locational shift toward the end of the midline primordium stage (18). However, we show here using more sensitive methods that *sim* (Vectorbase gene: AAEL011013) expression actually persists in the midline through to late midline stages, just like in other insects. Despite this commonality in expression pattern, *sim* transcriptional regulation at the midline primordium stages appears to have significant differences between the two species. While one *A. aegypti* enhancer shows remarkable conservation in position, composition, and mechanism with known *Drosophila sim* midline enhancers, the other is strikingly different from known *sim* enhancers in its location, time of onset, cell-type specificity, and lack of autoregulation. Our results suggest that a combination of conserved and novel regulatory mechanisms regulate the highly diverged *sim* locus of these two distantly-related dipterans (estimated divergence 241 MYA; (41)) to maintain a common gene expression pattern.

## RESULTS

### A. aegypti midline gene expression resembles that of other arthropods

We previously reported that *A. aegypti sim*, unlike its orthologs in other studied arthropods, shifted expression from the ventral midline to lateral regions of the embryonic CNS sometime during the midline primordium stage of midline development; this pattern was similarly adopted by other genes of the midline GRN functioning downstream of *sim* (18). We sought to establish a more thorough timeline for this shift in expression, as well as to determine more definitively whether *sim* and other midline genes were fully absent or merely severely reduced in expression in the late midline, using the sensitive hybridization chain reaction (HCR) method for in situ hybridization (42). Surprisingly, HCR revealed that *sim* expression remains strong in the *A. aegypti* ventral midline, with no shift to lateral CNS regions, through to late embryonic stages (Fig. 1A-F). Somatic muscle expression, similar to what has been observed in both *Drosophila* and in *Apis mellifera* (honeybee), was also observed (Fig. S1; (18, 24)). HCR probes for *short gastrulation* (*sog;* Vectorbase AAEL024210) and *shotgun* (*shg;* Vectorbase <u>AAEL012421</u>), two other genes previously reported to shift from midline to lateral in *A. aegypti* (18), similarly reveal a retained midline expression pattern (Fig. 1K-N).

**Figure 1:**
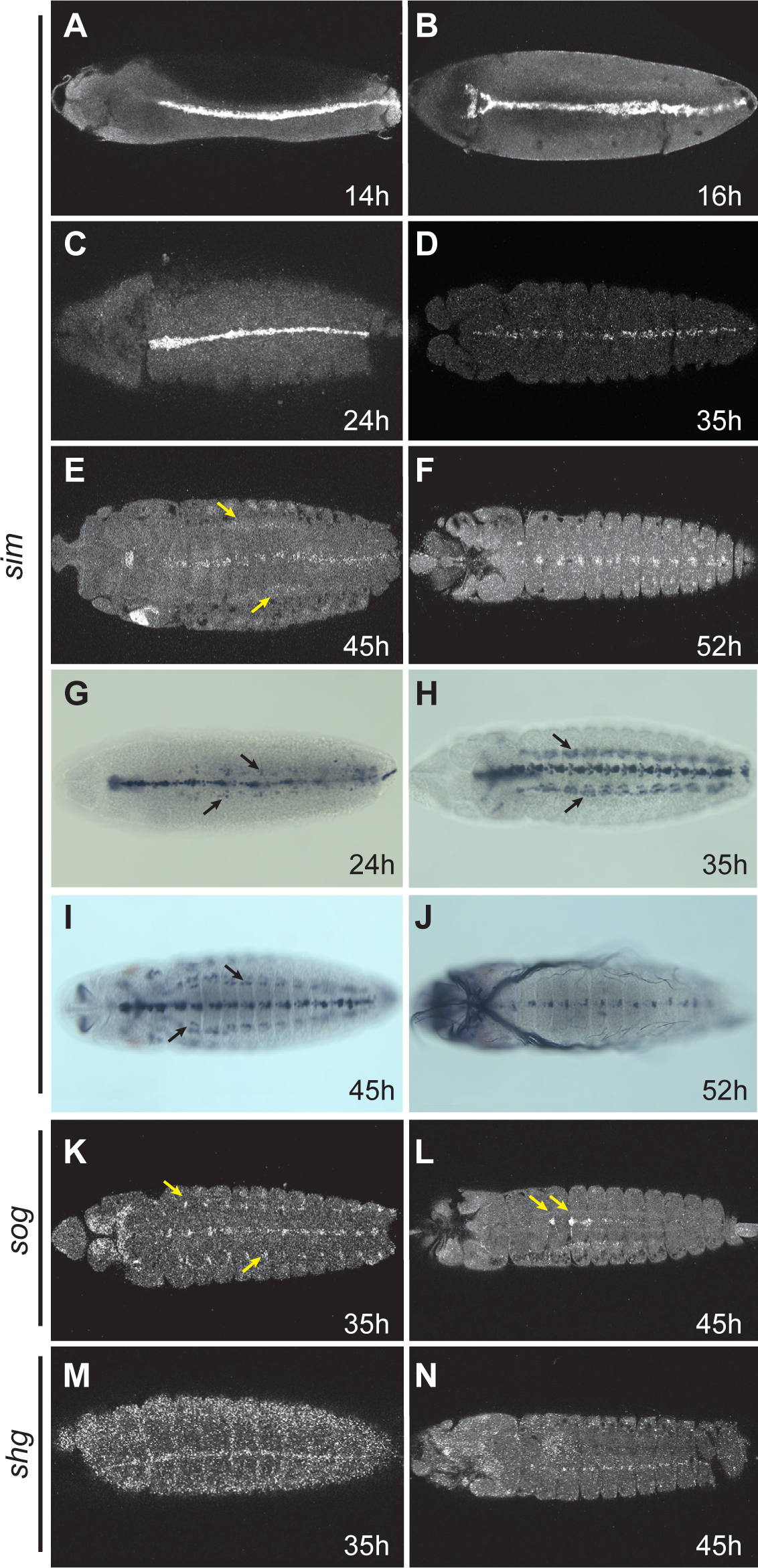
Expression of *sim*, *sog*, and *shg* in *A. aegypti* embryos. (A-F) Timecourse of *sim* expression in *A. aegypti* embryos from 14 hr. through 52 hr. using HCR. Note that expression is present in the midline at all stages. Lateral-appearing expression seen in panel E (arrows) is non-CNS expression in the somatic mesoderm (confirmed via *mef2* expression; see Supplemental Figure 1). (G-H) GFP expression from a *sim::GFP* fusion gene inserted in the native *sim* locus, visualized using anti-GFP antibody staining. Timepoints correspond to those in panels C-F. Arrows in G, H and I mark expression in the somatic musculature, not in the plane of focus in C and D (but see Fig. S1C,D). (K, L) *sog* expression in 35 and 45 hr. embryos as visualized by HCR. Limited expression is visible at the periphery of the CNS at 35 hr. (arrows); however, this is distinct from the more significant expression reported by Suryamohan et al. (18). Expression at 45 hr. is in cells whose morphology and position resemble the *Drosophila* dorsal median (DM) cells, which are not known to express *sog* and thus may represent a late embryonic expression difference between these two species. (M, N) Expression of *shg* as revealed by HCR in 35 and 45 hr. embryos. Note the prominent expression in the midline. In all panels, embryos are shown in their ventral aspect with anterior to the left.

To better understand the discrepancy between the current and previous results, we repeated the in situ hybridizations for *sim* and *sog* using the standard digoxygenin-labeled riboprobe/alkaline-phosphate detection protocol (43) on 45-hour *A. aegypti* embryos, with an extensive set of control experiments. We determined that the *A. aegypti* nervous system is largely refractory to this protocol during this mid-embryonic stage, yielding little true signal but a significant number of positive-appearing cells scattered throughout the tissue (Fig. S2). We detected little difference between sense and anti-sense probes (cf. panels E&H and F&I in Fig. S2), and obtained similar results with probes for the *Saccharomyces cerevisiae Gal4* gene, which has no mosquito homolog (Fig. S2D,G). The *sog* probe did have some limited activity in what appears to be a legitimate site of *A. aegypti sog* expression (analogous to the *Drosophila* dorsal median cells, Fig. S1J,K, arrows), but also ectopic-seeming expression elsewhere (Fig. S1K, arrowhead). In gut tissues, by contrast, we observed good overlap between the HCR and riboprobe methods (Fig. S2L, M, arrows), suggesting that the artifacts may be present primarily in nervous tissue (note the ectopic expression in the brain; Fig. S2L, M, arrowheads). Taken in sum, these results argue for greater reliability of the HCR data, and suggest that our previous reported results are incorrect.

To establish the *sim* expression pattern definitively, we used CRISPR/Cas9 genome editing to create a *sim::GFP* fusion gene in the *sim* locus (see Methods). As this fusion gene is under the control of the native *sim* genomic regulatory sequences, it should produce a faithful readout of *sim* expression. Sim::GFP was expressed in the midline and ventral somatic musculature identical to what we observed for *sim* HCR (Fig. 1G-J, cf. Fig 1C-F and Fig. S1C,D). Based on the HCR analysis and *sim::GFP* fusion data, therefore, we conclude that *A. aegypti* midline gene expression resembles that of *D. melanogaster* and other studied arthropods throughout embryogenesis.

### *Identification of* A. aegypti sim *enhancers*

Although the pattern of *sim* gene expression is similar between *D. melanogaster* and *A. aegypti*, we were curious to see what regulatory changes have occurred between the two species. The *Drosophila sim* locus is an order of magnitude smaller than its *A. aegypti* counterpart (Fig. S3) and, as is typical with insect species as many years divergent as *D. melanogaster* and *A. aegypti*, there is no readily detectable conservation of the non-coding sequences (5, 44). The *Drosophila* locus has been extensively assayed for regulatory sequences. At least four distinct enhancers are known to drive *sim* expression during the midline primordium stage, and one at the mesectoderm stage (Fig. S3B) (31, 35, 38). CRMs responsible for late midline expression remain unidentified. No *A. aegypti sim* enhancers are known. We chose eight initial sequences from the *A. aegypti sim* locus to assay for regulatory activity (Fig. S3A). Four were chosen based on sequence conservation with the related mosquito *Aedes albopictus* (estimated divergence 48 MYA; (41)), two were chosen as lying just upstream of the two annotated *sim* promoters, respectively, and two were selected essentially randomly from within the large first intron of the “B” transcript of the gene. As an initial screening assay, each selected sequence was tested for its ability to drive reporter gene expression in transgenic *Drosophila* embryos. Of the eight sequences, five had no observable expression (Fig. S3A), one drove expression in a pattern largely distinct from that of *sim* (Fig. S3A, Fig. S4), and two drove expression in the ventral midline (see below).

### *A mid-stage autoregulatory* sim *enhancer*

The *intP2* sequence is located in the *A. aegypti sim* first intron between the two annotated promoters and is roughly 40% conserved with *A. albopictus* (Fig. 2A). In *Drosophila* embryos, it drove reporter gene expression in a pattern identical to that of *sim* beginning at early stage 9 and persisting throughout embryogenesis (Fig. 2B-D). Closer examination of the sequence revealed that there are two primary blocks of sequence conservation (each about 75% identity) (Fig. 2A). We therefore tested each of these smaller conserved fragments individually. Whereas sequence *intP2A* failed to drive embryonic reporter gene expression (Fig. 2E), the 826 bp *intP2B* fragment has activity identical to that of the larger *intP2* sequence (Fig. 2F) and recapitulates endogenous *sim* expression (Fig. 2G). Consistent with a role as a *sim* enhancer, *intP2B* fully overlaps a region of open chromatin as assayed by FAIRE-seq (Fig. 2A; (45, 46)).

**Figure 2:**
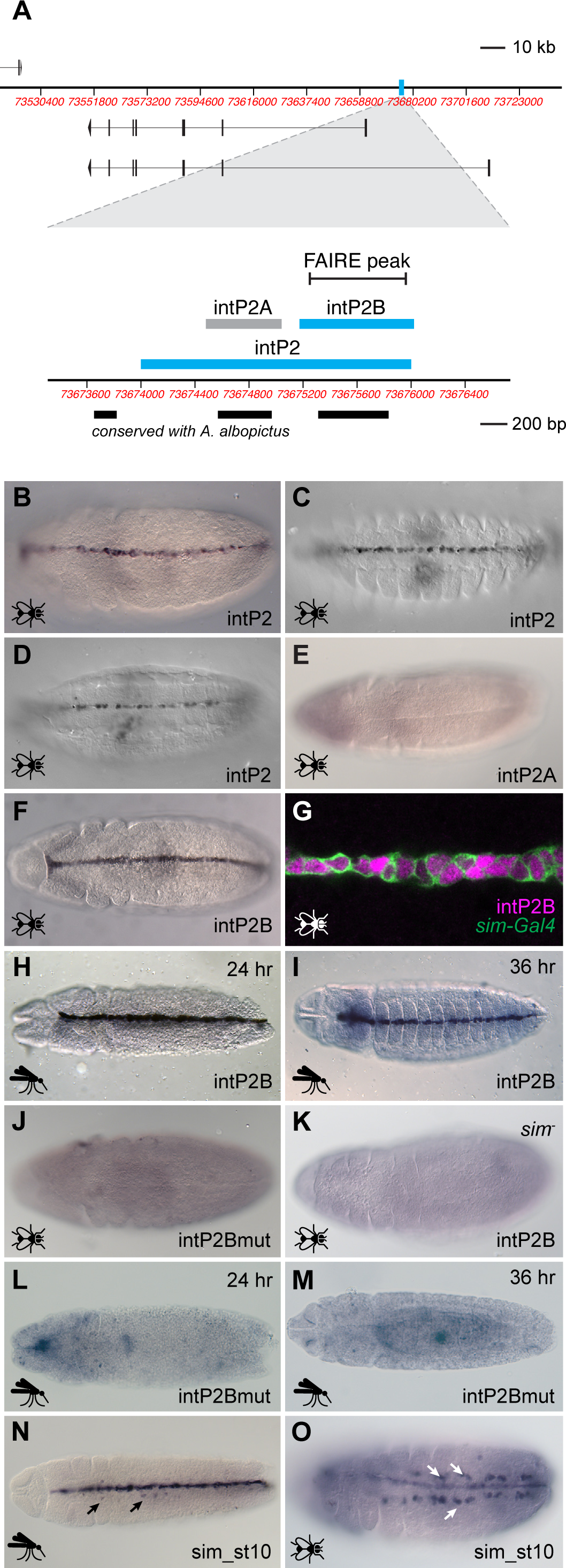
Characterization of the *intP2* enhancer. (A) Top: Schematic of the *A. aegypti sim* locus with the *intP2* sequence highlighted in cyan. Bottom: Close-up of the *intP2* region showing tested subfragments, conservation with *A. albopictus*, and FAIRE data. (B-O) Experiments in *Drosophila melanogaster* embryos are indicated by a fly icon, while those in *A. aegypti* embryos have a mosquito icon. Fly embryos are all stage 11, except for panels C (st. 14) and D (st. 15). All embryos are oriented ventrally with anterior to the left. (B) Reporter gene expression driven by *intP2* in transgenic *Drosophila* at stage 11, (C) at stage 14, and (D) at stage 15. (E) The *intP2A* fragment fails to drive reporter gene expression. (F) Reporter gene expression driven by *intP2B* in transgenic *Drosophila*. (G) Double-labeling confirms that *intP2B* activity (magenta) is in all midline cells, as marked by *sim-Gal4::UAS-tauGFP* (green). (H, I) Reporter gene activity driven by *intP2B* in 24 hr. (germ-band extended) and 36 hr. (germ-band retracted) *A. aegypti* embryos, respectively. (J) Mutation of CME sites in *intP2B* abrogates enhancer activity in transgenic *Drosophila*. (K) No *intP2B* activity is observed in a *sim* null background. (L, M) As in *Drosophila*, mutation of CME sites in *intP2B* abrogates enhancer activity in transgenic *A. aegypti*. (N) The *Drosophila sim_st10* enhancer drives reporter gene expression in the midline and in the ventral musculature (arrows) in *A. aegypti* transgenic embryos (cf. Fig. 1G). A 24 hr embryo is shown. (O) Similarly, the *sim_st10* enhancer drives expression in the midline and ventral musculature (arrows) in *Drosophila* embryos (cf. Fig. S1A).

To ensure that the activity we observed was not due to the sequence being assayed in a *Drosophila* background, we generated transgenic *A. aegypti* with *intP2* and *intP2B* driving a GFP reporter gene. Reporter gene expression in the transgenic mosquitoes was fully consistent both with the *Drosophila* transgenic results and with endogenous *A. aegypti sim* expression as revealed by HCR and *sim::GFP* (Fig. 2H, I and Fig. S5; cf. Fig. 2B and Fig. 1A-J). Reporter expression was fully present by 16 hours of development (early germ-band extended) and persisted past germ band retraction to at least 45 hours.

In *Drosophila*, *sim* expression at the midline primordium stages is autoregulated via binding of Sim:Tango (Tgo) heterodimers to RCGTG “CNS midline enhancer” (CME) sites (26, 31, 47). Inspection of the *intP2B* sequence revealed four of these potential autoregulatory sites (Fig. S6), mutagenesis of which abrogated enhancer activity in both transgenic *D. melanogaster* and *A. aegypti* (Fig. 2J, L, M and Fig. S7). Furthermore, *intP2B* was unable to drive reporter gene expression in *D. melanogaster* when recombined into a *sim* null background (Fig. 2K). Collectively, these results demonstrate that *A. aegypti* midline *sim* expression at the midline primordium stage is regulated through an autoregulatory enhancer, similar to what has been characterized in *D. melanogaster*.

The *intP2B* enhancer is remarkably similar to the *Drosophila sim_st10* enhancer (38), one of at least four *Drosophila* CRMs with reported midline primordium regulatory activity. Despite sharing no appreciable sequence conservation, the two enhancers are alike both in overall position within the *sim* first intron and in their complement of 3-4 Sim:Tgo “CME” autoregulatory sites (Fig. S6). Other putative binding sites are also in common, including those for Ventral veins lacking (Vvl) and Twist (Twi), although roles for these have not been tested empirically. We tested the *Drosophila sim_st10* enhancer in transgenic *A. aegypti* to determine if the fly enhancer was functionally equivalent to the mosquito enhancer. Surprisingly, we observed not only strong midline activity, but also activity in the Sim-positive muscle precursors (Fig. 2N). No such muscle activity was observed in either species with the *intP2* or *intP2B* enhancers. As this activity was not previously reported in *Drosophila* for *sim_st10* (38), we generated transgenic flies using the *sim_st10* sequence and assayed reporter gene expression at stages 11 and later. Consistent with what we observed in transgenic *A. aegypti*, clear muscle expression driven by *sim_st10* was observed in the *Drosophila* embryos (Fig. 2O). Therefore, despite substantial similarity in both position and binding site composition, *intP2B* and *sim_st10* do not have identical activity.

### A novel second midline enhancer

The *5P3* sequence is located in a conserved region in the *sim* 5’ intergenic region roughly 19.8 kb upstream of the *sim-RB* promoter (Fig. 3A). When assayed in *Drosophila* embryos, it drove expression in a subset of midline cells as well as in a variety of non-midline interneurons in more lateral regions of the CNS (Fig. 3B). By testing a series of smaller overlapping sequence fragments (Fig. 3A), we were able to separate the midline activity from the lateral activity (Fig. 3C-I). The lateral activity, which is ectopic with respect to native *sim* expression in both fly and mosquito, does not become apparent until germ band retraction at stage 13 and maps to at least two independent sequences, *5P3G* and *5P3B* (Fig. 2C, G). *5P3G* lies within the region of conservation with *A. albopictus*. The midline activity is contained within the 173 bp *5P3F* sequence, begins at late stage 9/stage 10, and persists through late embryogenesis (Fig. 3H, I). Testing in transgenic *A. aegypti* confirmed that *5P3* behaves in its native *trans* environment identical to how it behaves in *Drosophila*; expression is consistently seen in the midline by 24 hours of embryogenesis (mid germ-band extended stage) and persists through at least 45 hours (Fig. 3J, K; Fig. S5). Strikingly, we see lateral ectopic expression in the later stages, following germ band retraction (approximately 32 hr.), similar to what was observed in the *Drosophila* assay (Fig. 3K, arrows; Fig. S5).

**Figure 3:**
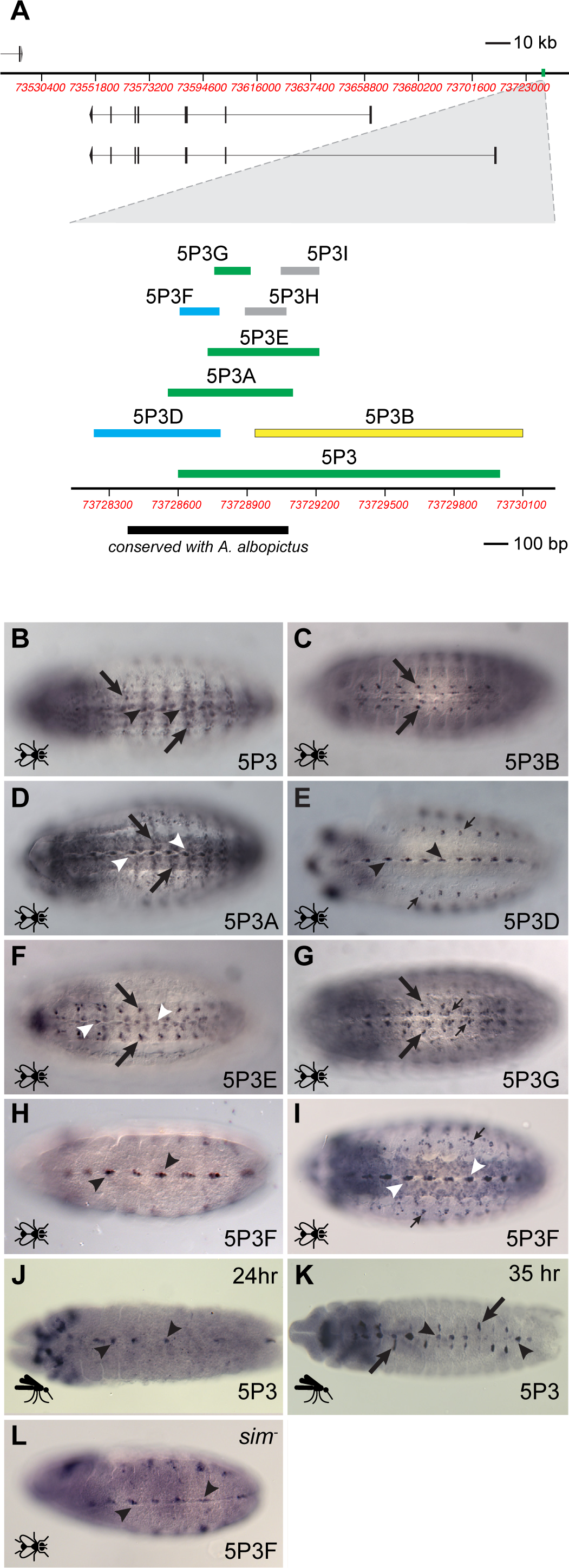
Characterization of *5P3* and its subfragments. (A) Top: Schematic of the *A. aegypti sim* locus with the *5P3* sequence highlighted in green. Bottom: Close-up of the *5P3* region showing tested subfragments and conservation with *A. albopictus*. Fragments with midline activity are colored cyan, those with lateral activity yellow, and those with both, green. Fragments with no enhancer activity are shown in gray. (B-K) Experiments in *Drosophila melanogaster* embryos are indicated by a fly icon, while those in *A. aegypti* embryos have a mosquito icon. Fly embryos are all stage 14-15 except for H, which is stage 11. All embryos are oriented ventrally with anterior to the left. (B) The *5P3* sequence drives reporter gene expression in midline (arrowheads) and lateral (arrows) CNS cells. (C) Activity of *5P3B* is restricted to lateral cells (arrows). (D) *5P3A* drives both midline (arrowheads) and lateral (arrows) reporter gene expression, whereas (E) *5P3D* is only active in midline cells (arrowheads). Non-CNS expression in the lateral bodywall (thin arrows) is a product of the expression vector, not the enhancer fragment. (F) *5P3E* drives both midline (arrowheads) and lateral (arrows) reporter gene expression, but (G) *5P3G* is confined to lateral regions only. Somewhat more medial expression (thin arrows) is not in the midline and is present, but out of the plane of focus, in the other fragments having lateral activity as well. (H, I) *5P3F* has midline expression only (arrowheads), pictured here at stages 11 and 15 respectively. Thin arrows in (I) indicate vector-induced expression. (J, K) The *5P3* fragment drives midline expression (arrowheads) in midline primordium-stage *A. aegypti* embryos, as well as ectopic expression following germ-band retraction (arrows). (L) Expression driven by *5P3F* is still observed in a *sim* null background (arrowheads), showing that *5P3F* is not subject to autoregulation.

Labeling using a variety of midline-expressed markers enabled us to identify the cells where the *5P3F* reporter gene is active in *Drosophila* embryos (Fig. 4). The stage 10 midline consists of 16 cells comprising three equivalence groups, the MP1, MP3, and MP4 groups (Fig. 4A; (48)). Subsequent signaling via the Notch pathway leads to specification of the anterior and posterior midline glia (AMG, PMG) and six neural precursors, MP1, MP3, MP4, MP5, MP6, and the median neuroblast (MNB). At late stage 11, these cells undergo Notch-mediated asymmetric cell divisions to form the MP1 neurons, the H-cell and H-cell sib neurons, and the Ventral Unpaired Median motorneurons and interneurons (mVUMs and iVUMs, respectively). The latter arise from MP4, MP5, and MP6, each of which divides to yield one mVUM and one iVUM. Specific combinations of gene expression allow each of these cells types to be uniquely identified (49). At stages 15-16, coexpression with Slit and Mab 22C10 (Futsch) revealed *5P3F* reporter gene activity in the posterior midline glia (PMG) and mVUMs, respectively (Fig. 4 C,D), while stage 13 expression just posterior to Nub expression rules out activity in the H and H-sib cells (Fig. 4B). At stages 13-14 we see coexpression with En, which marks the iVUMs, MNB and its progeny, and PMG, although reporter expression in PMG is inconsistent (Fig. 4E-H). At late stage 11, reporter expression is similarly coincident with En (Fig. 4I); the exception is the anteriormost cell (arrow), which we interpret to be one of the MNB progeny which loses En expression at about this stage (49). Taken together, these data suggest that the *5P3F* enhancer is specific to the early “MP4 equivalence group” (Fig. 4A, top), the earliest clear differentiation of the midline cells into specific fates (48).

**Figure 4:**
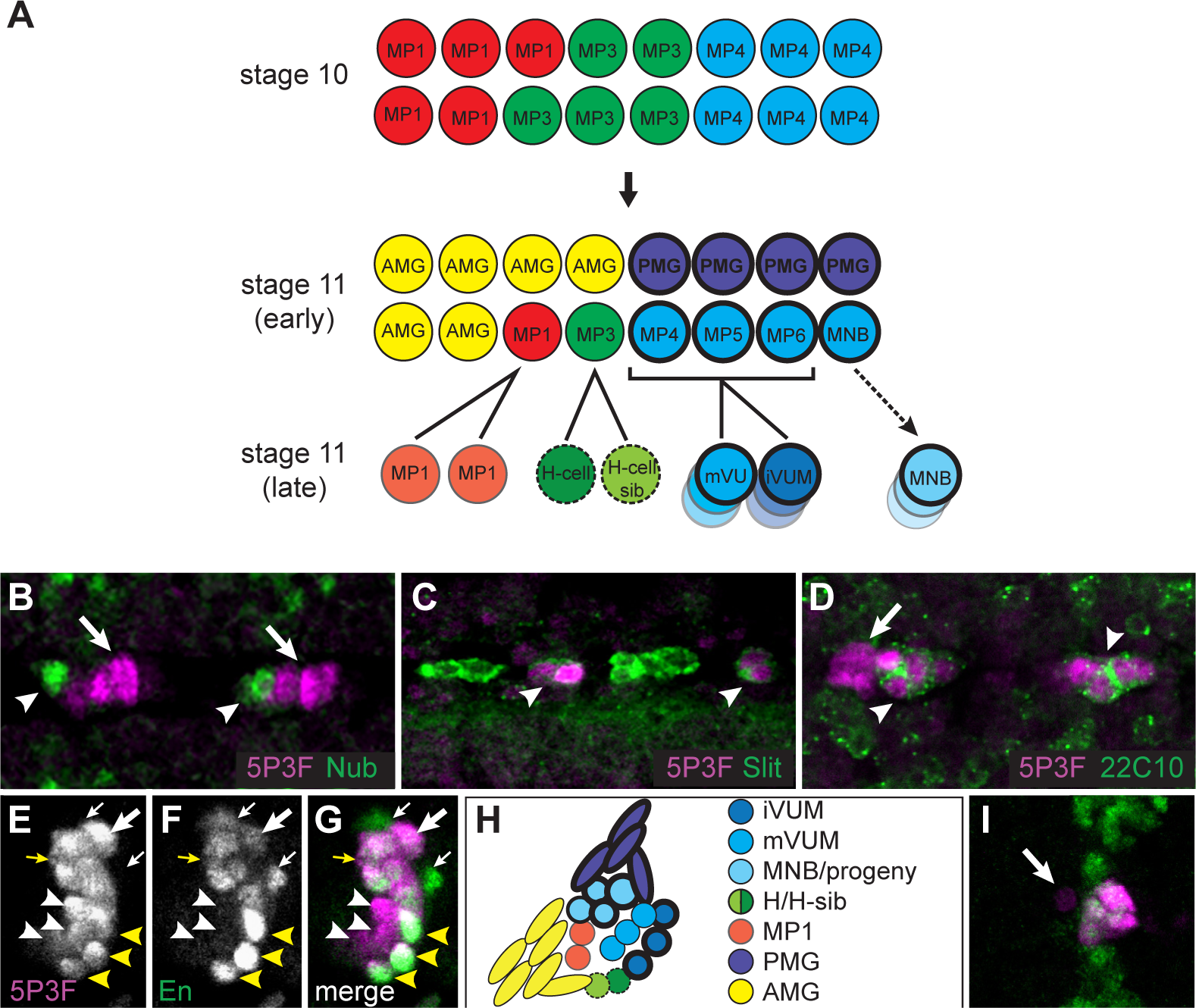
Cell-type activity of the *5P3F* enhancer in transgenic *Drosophila* embryos. (A) Cartoon of the midline cell lineages at the midline primordium stages (adapted from (68)). Cells of the MP4 equivalence group are shown in shades of blue. Cells with heavy outlining express Engrailed. (B) Co-labeling of the *5P3F* reporter (magenta) with Nubbin (Nub; green) at stage 13 shows that *5P3F* is not active in H-cell or H-cell-sib. (C) Co-labeling with Slit (Sli; green) indicates *5P3F* activity in posterior midline glia, shown in dissected embryos at stage 16. (D) *5P3F* activity is both in and adjacent to cells expressing Mab 22C10 (Futsch; green), which marks the mVUMs, shown here at stage 16. (E-G) Sagittal sections showing co-labeling of the *5P3F* reporter (magenta) with Engrailed (En; green) at stage 13. *5P3F* reporter gene expression is observed in the mVUMs (white arrowheads), iVUMs (yellow arrowheads), MNB and progeny (yellow arrow), and posterior midline glia (PMG; thick white arrow). Not all of the PMG are expressing the reporter (thin white arrows). (H) Cartoon of the stage 13 midline (adapted from (69)). (I) Co-labeling of the *5P3F* reporter (magenta) with Engrailed (En; green) at stage 11 (ventral view). All reporter-gene expressing cells are positive for En with the exception of a single anterior cell (arrow), likely one of the MNB progeny. Embryos in B, C, D, and I show a ventral section while E-G are sagittal sections; all embryos have anterior to the left.

Surprisingly, the *5P3F* sequence does not contain any *sim*-binding CME sites, suggesting that unlike all known *Drosophila* midline primordium-stage *sim* enhancers, it is not autoregulated. Consistent with this, reporter gene expression is maintained even in a *sim* null background (Fig. 3L). The *A. aegypti 5P3F* enhancer is therefore novel in several distinct ways as compared to both the *intP2* enhancer and to any of the known *Drosophila sim* enhancers: its activity begins later, at the stage 9/10 boundary; it is expressed in only a subset of the midline cells; it lacks autoregulatory capability; and it is located substantially upstream of the transcription start site.

### *Functional assessment of conserved regions from* Aedes albopictus

The *intP2* and *5P3* sequences were chosen as candidate enhancers in part due to sequence conservation with the related Aedine mosquito *A. albopictus*. We were therefore curious to determine whether these partly conserved *A. albopictus* sequences were also able to function as midline enhancers. For *intP2*, we selected a 648 bp sequence *A. albopictus* sequence similar to the *A. aegypti intP2B* fragment (Fig. 2A). The two sequences have 68% identity over a central 600 bp of alignment, including conservation of all four Sim:Tgo binding sites. For the *5P3* region, we chose the primary conserved region (∼70% identical) that aligns from the middle of the *5P3D* fragment through the *5P3H* fragment (Fig. 3A). Both of these *A. albopictus* fragments drove reporter gene expression in transgenic *D. melanogaster* in a pattern similar to their *A. aegypti* counterparts (Fig. S8).

## DISCUSSION

The *sim* locus in *A. aegypti* is an order of magnitude larger than its well-studied counterpart in *D. melanogaster*, with major increases in the lengths of both introns and the 5’ intergenic region. Moreover, the non-coding sequences of the two loci have diverged well past the point of detectable similarity. We show here that despite this dramatic expansion and divergence, there exists a striking conservation of core aspects of *sim* regulation. At the same time, apparently novel regulatory mechanisms are also observed.

### Conserved and novel enhancers

Although the *intP2B* enhancer is remarkably similar to the *Drosophila sim_st10* enhancer, it lacks the muscle activity regulated by the latter. Several possibilities may account for the differences in activity: the *sim_st10* sequence may be a composite of two enhancers, a midline enhancer and a muscle enhancer, and *intP2B* is the counterpart of just the midline one; the proper counterpart to *int2B* in *Drosophila* may be one of the other three or more identified *sim* midline enhancers (none of which have described muscle activity), despite their being less similar to *int2B* in terms of position and binding site composition; or the sequence may have evolved different capabilities in the two species. Future finer-scale mapping of both the *A. aegypti* and *Drosophila* loci, with identification of additional enhancers and further dissection of known ones, will be necessary to distinguish between these scenarios.

The *5P3F* enhancer, by contrast, is completely novel compared to known *Drosophila sim* enhancers with respect to its spatial activity, temporal activity, and lack of autoregulatory capability. The latter is particularly striking, as all studied *Drosophila* midline enhancers contain CME sites (25–31). Full activity of all tested midline enhancers requires functional CMEs, although in a minority of cases reduced enhancer activity can still be observed in their absence (25). Hong et al. (31) have noted a correlation between number of CME sites and onset of enhancer activity, with a high number of binding sites (four) needed for early midline primordium activity. Although *5P3F* has a later onset than *intP2B* and the known *Drosophila sim* enhancers, its activity by stage 10 still places it in the class requiring at least two CME sites by their analysis. *5P3F* function, therefore, is likely to be governed by novel regulatory mechanisms.

*5P3F* is also unique in that it appears to initiate activity only in the MP4 equivalence group. All currently known *Drosophila sim* enhancers that initiate at the midline primordium stage do so in the complete set of midline cells. *5P3F* activity is most like that of the *mfas_780* enhancer (25), which is active in the MP4-6 cells as well as in anterior midline glia. *5P3F*, however, only drives expression in the posterior, not anterior, midline glia. When tested in *Drosophila* embryos, *5P3F* reporter gene expression is similar to that of En at stages 10-11; later deviations may simply result from perdurance of the *lacZ* reporter in cells where En has already ceased expression.

Further studies will be necessary to determine the off times of both the *intP2B* and *5P3F* enhancers. Interestingly, despite the extensive characterization of the *Drosophila sim* locus, enhancers active in late midline stages have not been isolated. Thus, sequences regulating *sim* expression following germ band retraction, if distinct from those active at the midline primordium stage, remain unknown.

Importantly, it is also not currently known if the cellular makeup of the *A. aegypti* midline is one-for-one identical to that of *Drosophila* or not, and if gene expression patterns are completely conserved. For example, while the overall morphology, number, and arrangement of neuroblasts in the beetle *Tribolium castaneum* is conserved with *Drosophila*, there is substantial variation in gene expression patterns in the neuronal lineages (50). Thus, extensive additional analysis of the *A. aegypti* CNS remains necessary in order to put our results fully in context. The reporter lines we have generated will be of considerable aid in this regard.

While the regulatory mechanisms governing *5P3F* are novel compared to known *Drosophila sim* enhancers, we note that *5P3* reporter gene activity is identical in all respects when tested in both *A. aegypti* and *D. melanogaster* transgenic embryos. This includes its later onset of expression at the mid, not early, midline primordium stage and presence of ectopic activity in lateral CNS cells following germ band retraction. Therefore, the same regulatory mechanisms are available for use in both species. It will be interesting to determine whether a *5P3F*-like enhancer exists in the *Drosophila sim* locus, or if the *A. aegypti* sequence is a novel regulatory element that is tapping into an available, but not accessed, regulatory capability. Kalay et al. (15) have described “cryptic enhancers,” sequences with regulatory capability that are repressed by surrounding sequences in some species, but which constitute latent enhancers that could potentially be activated in other species to evolve a new expression pattern. Further characterization of the *Drosophila sim* locus, including finer-scale dissection of known enhancer regions, will be necessary to establish whether this is the case, whether there is an active but as-yet unidentified MP4-equivalence group *sim* enhancer, or whether the exogenous *5P3F* sequence is independently responding to signals in the *Drosophila trans* environment without having a *Drosophila* genomic counterpart.

We used here *Drosophila* transgenic reporter lines as an expedient method to test our putative enhancers for activity, and only then tested the best candidates in their native *A. aegypti* host. This is a common strategy that has served well in the past. However, it will still be necessary at some point to confirm whether the sequences that failed in the *Drosophila* assay also fail in their native *A. aegypti trans* environments. This is especially true for the *intP3* sequence, which was able to drive expression in a non-*sim* like pattern in *Drosophila*.

### Challenges of non-model organisms

Special challenges come into play when studying regulatory evolution in non-traditional laboratory species (51) and at large evolutionary distances. Fortunately, *A. aegypti* has proven a tractable experimental system, with robust transgenic capability and straightforward husbandry. Nevertheless, our experiences underscore the care that must be taken when working in relatively unexplored territory, as “standard” techniques such as digoxygenin/riboprobe based in situ hybridization may prove unreliable, and orthogonal methods and corroborating data are not always available. In this case, use of the sensitive HCR method plus an extensive set of control experiments, along with generation of an engineered GFP fusion gene, enabled us to determine that previous characterizations of the *A. aegypti sim* expression pattern were incorrect, and allowed for a convincing updated description. However, other challenges remain. As mentioned above, detailed morphological and molecular-level investigations of the *A. aegypti* CNS are still required to properly establish homology with *Drosophila* between the midline and other nervous system cell types. Moreover, the high degree of non-coding sequence divergence between the two species makes identifying regulatory sequences a particular challenge. While we demonstrate some success here by using conservation between the two Aedine species *A. aegypti* and *A. albopictus*, sequence alignment in general provides a limited means for enhancer discovery (52). The recent availability of additional Aedine genomes (53) may be of future help, as will success we have had using computational methods such as SCRMshaw to identify functionally-related regulatory sequences across the holometabolous insects (5, 18, 54, 55).

### Regulatory evolution of a locus

Enhancer activity within a locus can be maintained over extraordinary evolutionary distances. For example, sequences from the sponge *Amphimedon queenslandica Islet* locus are able to drive reporter gene expression in transgenic zebrafish in patterns reminiscent of (although not identical to) the zebrafish *Islet2a* gene (7), and in similar tissues in transgenic mice, despite over 750 MY of divergence. Similarly, a deeply conserved region of the *SoxB2* locus maintains related although not identical function in species from vertebrates to cnidarians (∼700 MY). Strikingly, both human (6–8) and sea urchin versions of this enhancer are also able to drive reporter gene expression in a *Sox*-related pattern in *Drosophila*, despite no obvious sequence homology (6). While these studies make clear the remarkable ability of enhancers to retain functional conservation over hundreds of millions of years, the vast distances and limited characterization of the respective loci make it difficult to infer how the complete regulatory locus may have evolved.

The overall regulatory evolution of a single locus has previously been evaluated extensively in the case of the *Drosophila yellow (y;* FlyBase FBgn0004034) gene, which encodes a protein in the pigmentation pathway. Kalay et al. (12) found that enhancers with similar tissue-specific activity in three *Drosophila* species were often found in different locations within their respective loci. However, the authors note that *y* expression is a highly evolving trait, and speculate that enhancers controlling conserved aspects of expression may be more constrained in genomic location. The analysis is complicated by the subsequent identification of numerous redundant and semi-redundant enhancers in the *y* locus (15). Determining which enhancers have changed location versus which have maintained position is thus a non-trivial exercise.

Frankel et al. (56) undertook a thorough analysis of the *shavenbaby* (*svb*; FlyBase: *ovo,* FBgn0003028) locus in *D. melanogaster* and its 30-40 MYA diverged relative *D. virilis*. In the case of *svb*, which encodes a transcription factor that serves as a “master regulator” of trichome development, the position and overall function of enhancers appears to be maintained. Although like pigmentation trichome patterning is a rapidly-evolving trait, *svb’s* more pleiotropic role as a master transcriptional regulator may induce stronger constraints on the overall regulatory architecture of its locus (consistent with the speculations of Kalay et al. (12) discussed above). Enhancer positioning has also been explored in several cases of much larger evolutionary divergence than the 25-50 MYA range examined for *y* and *svb* regulation. Cande et al. (4) looked at the locations of enhancers for several developmental genes in species as diverse as *D. melanogaster, T. castaneum, Anopheles gambiae* (mosquito), and *Apis mellifera* (honeybee). Many of these enhancers appear to be maintained in a similar genomic position even over these large distances (up to 345 MYA), although the locations of others seem not to have been conserved. Similar results were observed by Kazemian et al. (5). These analyses are complicated by the fact the overall regulatory loci are not defined in detail, making it difficult to assess the full regulatory landscape in the way that was done for *y* and *svb*, albeit among more closely-related species.

The *Drosophila sim* locus, by contrast, has been characterized in detail, with almost all non-coding sequences in the ∼30 kb locus tested for enhancer activity. *sim* is a core developmental gene with embryonic midline activity conserved over more than 350 MY of arthropod evolution. Our initial analysis of *A. aegypti sim* enhancers suggests both conserved position and mechanism of action for some enhancers, but also potentially novel regulatory mechanisms acting through non-homologous enhancers at new locations. Further studies of this locus will be necessary to determine whether or not there are additional novel regulatory activities, and to what degree the various midline as well as non-midline enhancers are retained, lost, or altered. It has been well-established that developmental genes often have highly conserved expression patterns mediated by enhancers with common sequence characteristics, despite these enhancers having frequently diverged past the point of recognition via simple sequence comparisons. Unresolved, however, is the question of whether or to what extent such functionally-equivalent enhancers are conserved by direct descent or result from convergent processes (5). Understanding how the genomic organization of these enhancers changes over evolutionary time aids in clarifying this question. Although an analysis of only two species, such as that presented here, cannot answer the descent vs. convergence question, the observation that at least some enhancers appear to be maintained in analogous locations suggests one of two possibilities: the enhancer has been conserved through direct descent, or there is a functional constraint on the positioning of a regulatory element with the necessary function, leading to convergence of not just sequence composition but also of positioning for a newly-evolved enhancer. Although the latter seems to us the less probable alternative, it raises intriguing questions about locus-level regulation of gene expression. While the “grammar” of individual regulatory elements has often been discussed (57), potential larger-scale grammars have received much less attention. Whether the arrangement, spacing, or other aspects of the relative configuration of the various enhancers in a locus has regulatory consequences remains an open and largely unexplored frontier. Continued analysis of the well-explored and tractable *sim* locus should help to shed light on this and other important questions relating to how the regulatory landscape of a major and highly-conserved developmental regulator evolves.

## Materials and Methods

### Reporter constructs and transgenic Drosophila

Sequences were amplified by PCR on genomic DNA isolated from *A. aegypti* Liverpool (LVP) wildtype mosquitoes. Eighteen putative *sim* enhancer sequences were cloned, using the Gibson Assembly method (58), into plasmid *pLacZattB*, a φC31-enabled *Drosophila* transformation vector containing the *LacZ* gene under the control of a minimal *hsp70* promoter (59). Primer sequences are provided in Table S1. Transgenic flies were produced by Rainbow Transgenic Flies (Camarillo, CA) by injection into line attP2 8622.

### *Reporter constructs and transgenic* A. aegypti

Sequences were amplified by PCR on genomic DNA isolated from *A. aegypti* LVP wildtype mosquitoes, with the exception of *sim_st10*, which was synthesized as a gBlock (IDT, Coralville IA) based on *D. melanogaster* dm6 sequence 3R:13060273..13060969. The *intP2*, *5P3,* and *sim_st10* sequences were cloned into pENTR-D-Topo (Thermo Fisher) and then by Gateway recombination (60) into *pgPhiGUE*, a *piggyBac* transformation vector containing enhanced green fluorescent protein (EGFP) under the control of the *Drosophila* synthetic core promoter (gift of Y. Tomoyasu, Miami University). Primer sequences are provided in Table S1. Transgenic mosquitoes for *intP2* and *5P3* were produced by the Insect Transformation Facility at the Institute for Bioscience & Biotechnology Research (Rockville, MD) by injecting preblastodermal *A. aegypti* LVP embryos; the other transgenic mosquito lines were generated in the Halfon lab. Genomic insertion sites for the transgenic lines were identified using the Splinkerette method (61) and are provided in Table S2. Control lines consisting of the *pgPhiGUE* vector with a non-regulatory dummy sequence inserted in lieu of an enhancer confirmed that there is no vector-dependent embryonic reporter gene activity (Fig. S9).

### *Construction of the* sim::eGFP *fusion line of* A. aegypti

An active sgRNA within Exon VII of the *sim* locus was identified by microinjection of *A. aegypti* LVP embryos co-injected with 80 fmol/µL each of the *pHsp70-Cas9* vector (Addgene 45945) and an sgRNA expressing plasmid containing the U6:3 promoter for AAEL017774 (U6 spliceosomal RNA) followed by the sgRNA scaffold from (62) programmed for the protospacer target site. The active sgRNA site was identified 39 bp upstream of the *sim* stop codon at position Chr1:73549491-73549513. A homology arm donor was then synthesized as a gBlock (IDT, Coralville IA) with the donor cargo flanked by 400 nucleotides up- and downstream of the CRISPR/Cas9 cleavage site (Table S3). The donor consisted of a synonymous SNP recoding of the *sim* 39 C-terminal nucleotides, a 39 nucleotide fusion linker terminating with an EcoRI cut site followed by five additional nucleotides, and an XhoI restriction site. The U6:3 promoter for AAEL017774 followed by the sgRNA scaffold from (62) programmed for the active cut site in sim Exon VII was further included in the gBlock, downstream of the right homology arm (see Fig. S10). The gBlock was cloned into pBluescript using ApaI and NheI/XbaI. The eGFP fusion, terminator, and *3xP3-dsRed* marker were then amplified from the pgPhiGUE vector using primers eGFP-fus-EcoRI-F and 3xP3-fus-XhoI-R (Table S3), and cloned into the intermediate plasmid using EcoRI and XhoI. Lines were generated in the Halfon lab where preblastoderm embryos of *A. aegypti* LVP were microinjected with 80 fmol/µL each of the donor and the pHsp70-Cas9 helper. Surviving G0 animals were backcrossed to wildtype LVP and the G1 progeny were screened for the presence of the *3xP3-dsRed* eye marker. Correct insertion of the donor construct was confirmed by Nanopore sequencing of a PCR product using primers Aeg_sim_E1_F1 and 3xP3-Forward, which spans the 3xP3 marker and runs into the *sim* gene beyond the region included for the homologous recombination (Figure S8).

### Mosquito husbandry

This project used the *A. aegypti* LVP_ib12 strain, obtained from the Malaria Research and Reference Reagent Center (MR4; BEI Resources). *A. aegypti* lines were reared in an insectary at constant conditions of 27 °C, 80% relative humidity, and a 12:12 hr light:dark cycle. Larvae were reared in pans and fed on ground Tetramin fish flakes (Spectrum Brands Pets, Blacksburg, VA). Adult mosquito diet was raisins, with adult females fed three times every five days with defibrinated sheep blood (Hardy Diagnostics).

### Immunohistochemistry

For all analyses, a minimum of ten embryos were analyzed in detail and for transgenic mosquitoes, at least four independent insertion lines were examined. *Drosophila* embryo preparation and immunohistochemistry was performed using standard methods. *A. aegypti* embryo preparation and staining was performed essentially as described by Clemons et al. (63, 64) with modifications as detailed in Reid et al. (65). Primary antibodies used were mouse anti-β-galactosidase (1:500; Ab-cam), rabbit anti-GFP (1:500; Ab-cam, ab290), rabbit anti-β-galactosidase (1:500; Ab-cam), mouse anti-Slit C555.6D (1:10; Developmental Studies Hybridoma Bank), mouse anti-Nubbin 2D4 (1:10, DSHB), mouse anti-En 4D9 (DSHB), and Mab 22C10 (1:20, DSHB). The ABC kit (Vector Laboratories) was used for immunohistochemical staining. For fluorescent staining, the secondary antibodies used were anti-mouse Alexa Fluor 488 (Molecular Probes, A11029; 1:250) and anti-rabbit Alexa Fluor 647 (Molecular Probes, A21245; 1:500). Fluorescent staining was visualized by confocal microscopy using a Leica SP8 confocal microscope.

### In situ *hybridization*

*A. aegypti* embryos were prepared as described by Reid et al. (65). In situ hybridization using digoxygenin-labeled riboprobes was performed as in Haugen et al. (43). For HCR, reagents were obtained from Molecular Instruments (Los Angeles, CA). Embryos were stained as described by Bruce et al. (66) with the following modifications (67): Embryos were incubated at 37°C overnight with HCR probes at a concentration of 16 µL/mL, washed 5x in probe wash buffer, then incubated overnight in the dark in 100µL of a 6% solution (each) of the labeled HCR hairpins. Embryos were subsequently washed three times in 5xSCCT, one time with 1xPBS, then resuspended in 80% glycerol containing 4% n-propyl gallate and visualized by confocal microscopy using a Leica SP8 confocal microscope.

### Motif Analysis and Mutations

The *intP2B* enhancer sequence was scanned for exact matches to the Sim binding site consensus sequences ACGTG (47) and DDRCGTG (31) using MacVector (Apex, NC). Four sites were identified, and each motif was mutated to the BamHI restriction site GGATCC as in Wharton et al. (26). The mutated enhancer sequence was synthesized as a gBlocks Gene Fragment (IDT, Coralville IA) and inserted into *pattBnucGFPh* using the NEBuilder HiFi DNA Assembly kit (New England Biolabs, Beverley, MA).

### Test for autoregulation

The *intP2B* and *5P3F* reporter constructs were recombined onto a *sim* null mutant chromosome (*sim^2^*, Bloomington Stock Center stock #2055) and balanced over a *ftz-lacZ* containing *TM3* balancer.

### Image credits

The following graphical elements were obtained from The Noun Project (thenounproject.com) under a Creative Commons CC-BY 3.0 licence: fly, Georgiana Ionescu; mosquito, Cristiano Zoucas.

## ACKNOWLEDGMENTS

We thank Katie Costanzo and Molly Duman-Scheel for help and advice on mosquito rearing, Yoshi Tomoyasu for the pgPhiGUE plasmid and for helpful comments on the manuscript, Jack Leatherbarrow for assistance with staining and cloning, Cassandra Rorhdanz for help with mosquito husbandry, and members of the Halfon lab for useful advice and discussion. Fly stocks obtained from the Bloomington Drosophila Stock Center (NIH P40OD018537) were used in this study. Monoclonal antibodies were obtained from the Developmental Studies Hybridoma Bank, created by the NICHD of the NIH and maintained at The University of Iowa, Department of Biology, Iowa City, IA 52242. This study also utilized the resources of the Jacobs School of Medicine and Biomedical Sciences Confocal Microscopy Core Facility.

## SUPPLEMENTAL LEGENDS AND TABLES

**Supplemental Table S1:**
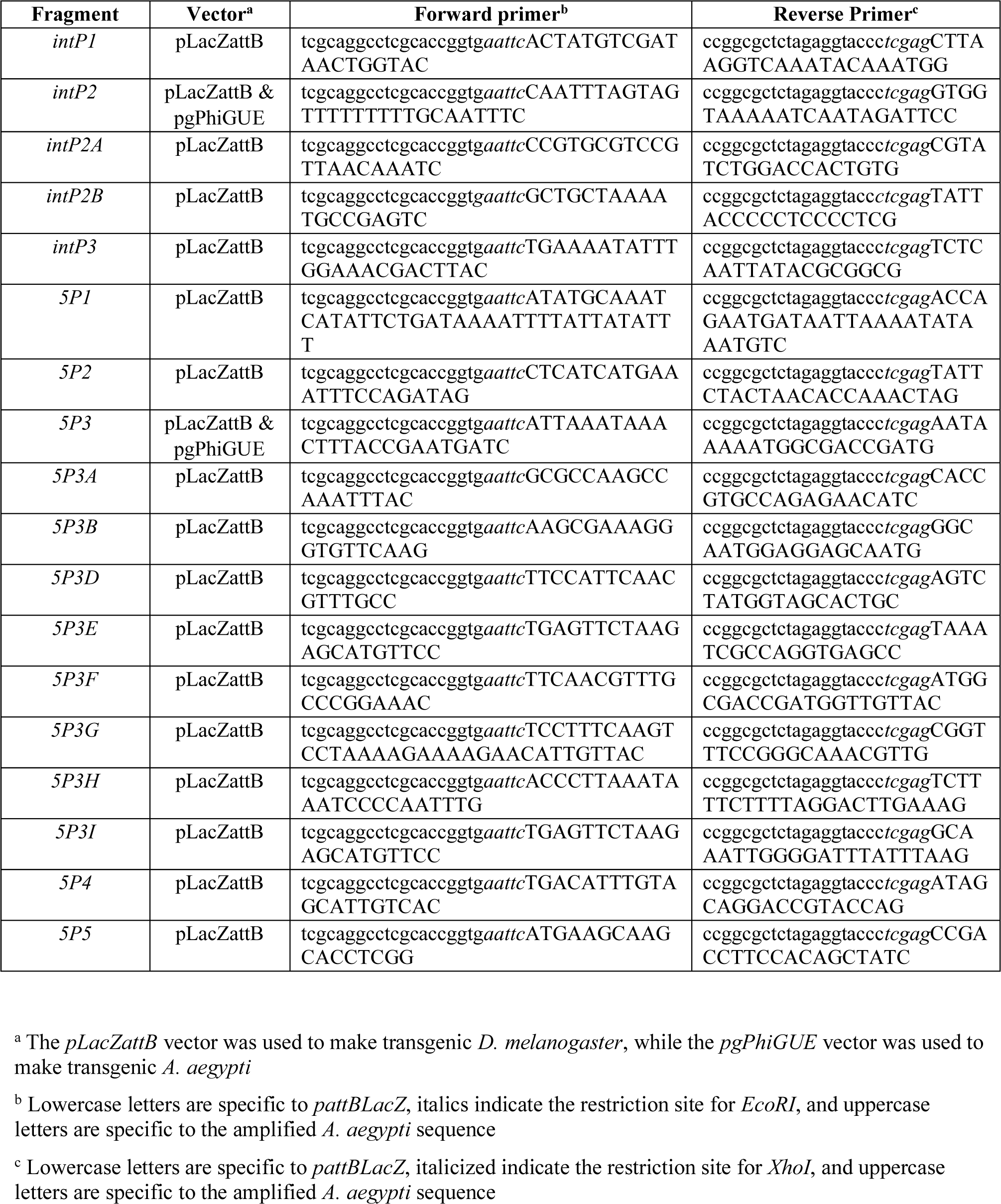
Primers for transgenic fly and mosquito lines

**Supplemental Table S2.**
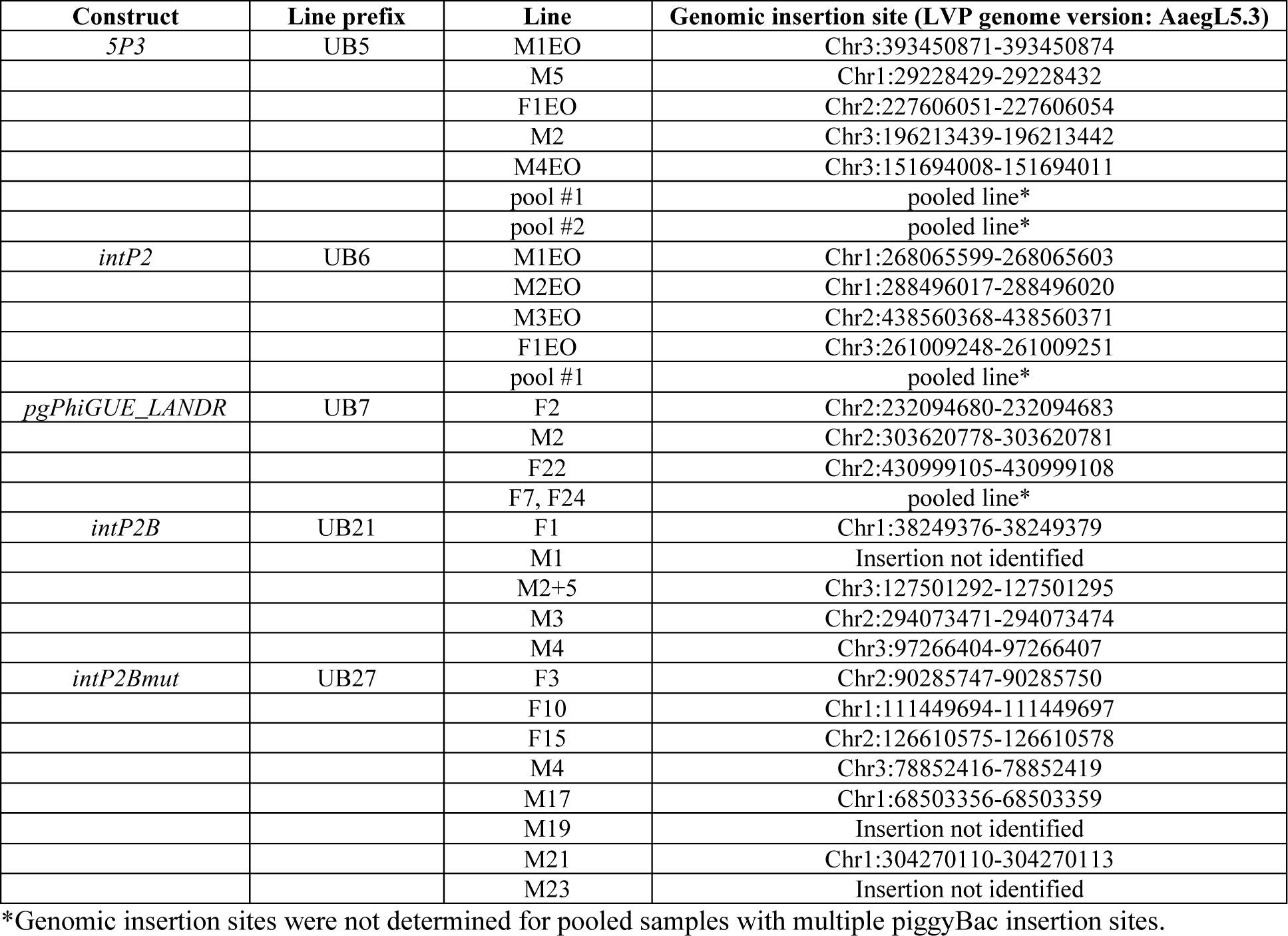
Genomic insertion sites for the transgenic *A. aegypti* lines

**Supplemental Table S3.**
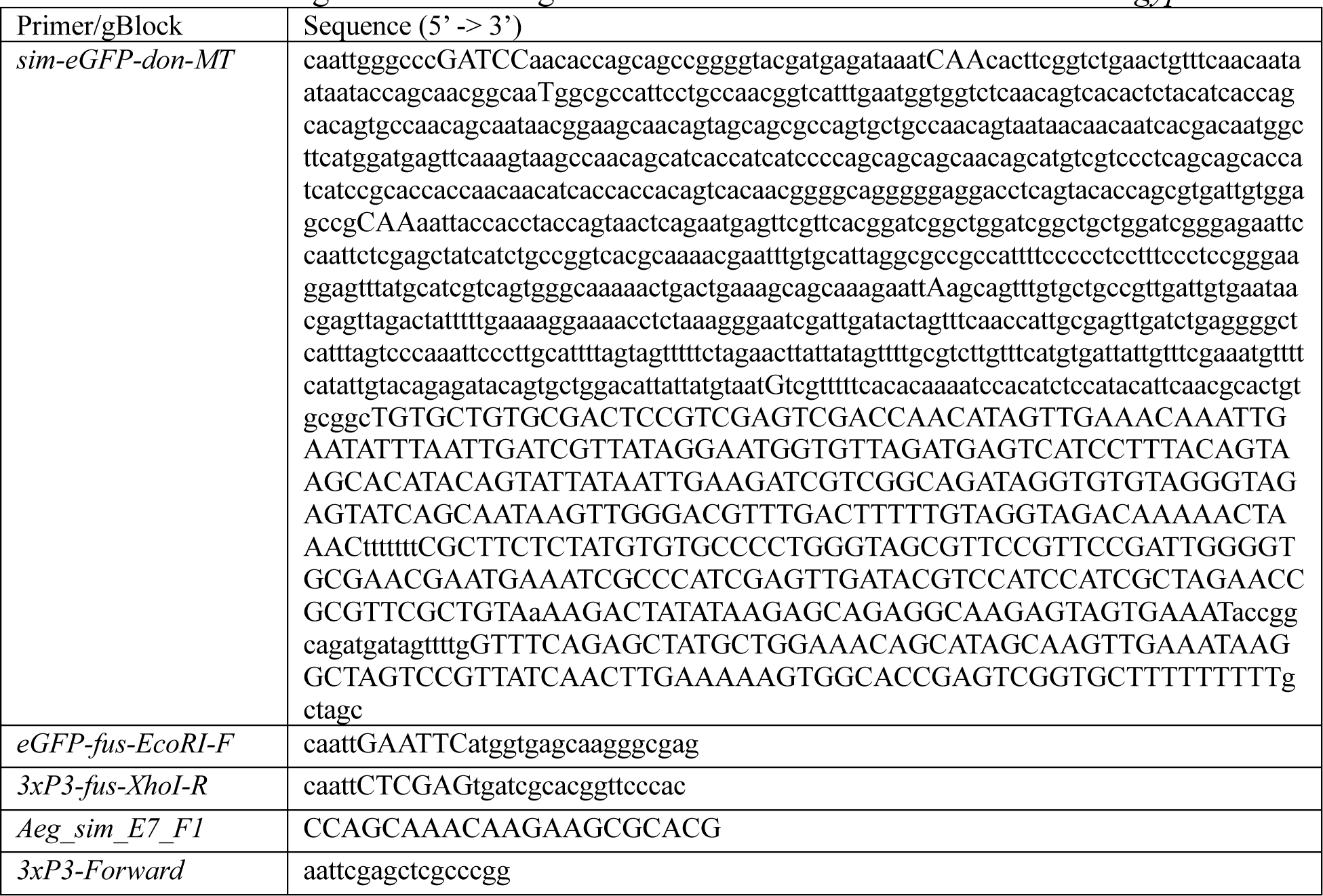
Primers and gBlocks used to generate the *sim::eGFP* fusion line of *A. aegypti*.

**Supplemental Figure S1:**
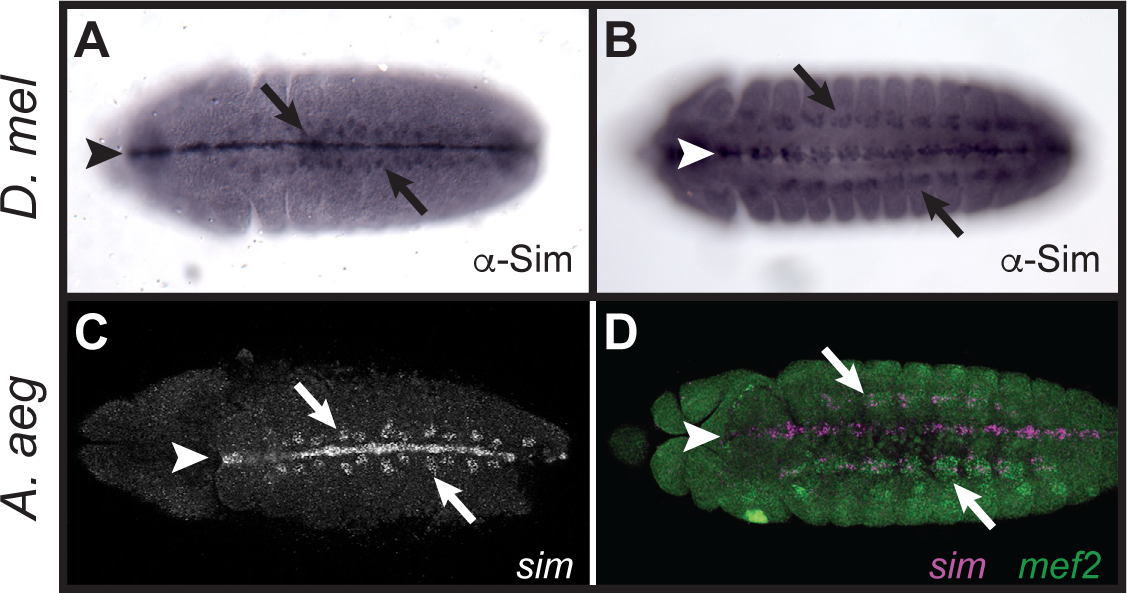
*A. aegypti sim* is expressed in muscle precursors and developing somatic muscle. (A, B) *Drosophila* embryos stained with anti-Sim antibodies at stages 11 and 14, respectively, show Sim expression in both the midline (arrowheads) and muscle (arrows). (C, D) Similar expression is detected by HCR in *A. aegypti* embryos at analogous stages. The embryo in D is co-labeled with HCR for *sim* (magenta) and the muscle-specific gene *mef2* (green).

**Supplemental Figure S2:**
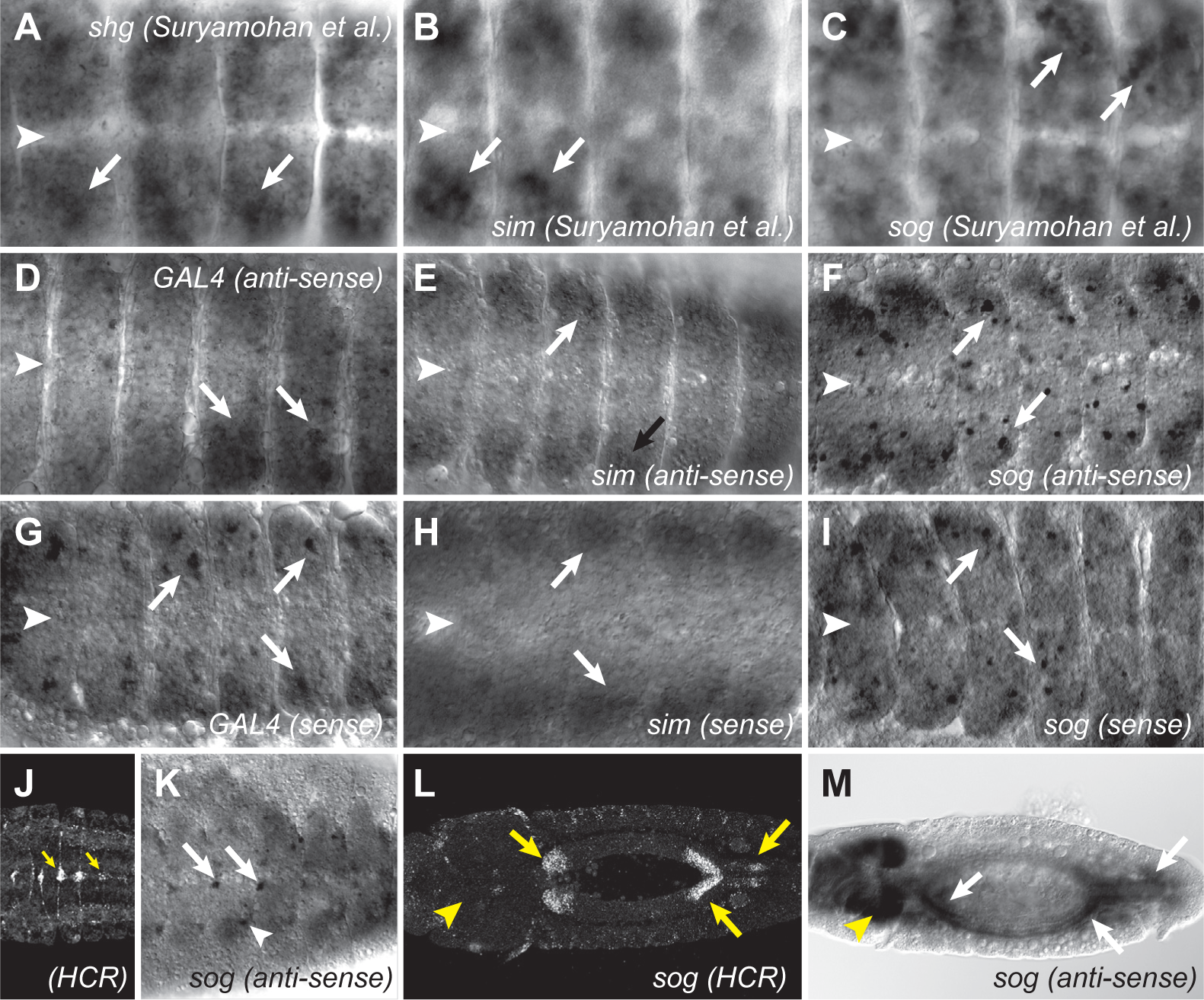
In situ hybridization artifacts in the *A. aegypti* embryonic nervous system explain earlier erroneous expression pattern data. (A-C) In situ hybridization results reported by Suryamohan et al. (18) for the *A. aegypti* orthologs of *Drosophila* (A) *shg*, (B) *sim,* and (C) *sog*. All three genes appear to be absent from the midline (arrowheads) but expressed in lateral regions of the nerve cord (arrows). (D-F) New in situ hybridization results using anti-sense probes for yeast *Gal4* (D), and *A. aegypti sim* (E) and *sog* (F). Note again the overall absence of expression in the midline (arrowheads) and presence in lateral nerve cord regions (arrows). (G-I) In situ hybridization using control (sense) probes for the same sequences as D-F. Note the qualitatively similar appearance of limited midline but prominent lateral expression. (J, L) Results for HCR against *A. aegypti sog* compared to (K, M) standard in situ hybridization results suggest that the standard in situ method does work in some tissues, but is heavily artifact prone in others. Although among the positive-appearing cells in panel K are midline cells morphologically similar to the dorsal median cells of *Drosophila* (arrows), which are clearly positive by HCR in J (arrows), there are also lateral cells (arrowhead) with no corresponding HCR signal. Similar positive signal can be observed in the anterior and posterior midgut in L and M (arrows), but note the apparent artifact of strong signal in the brain in M whereas there is no corresponding expression observed by HCR in L (arrowheads).

**Supplemental Figure S3:**
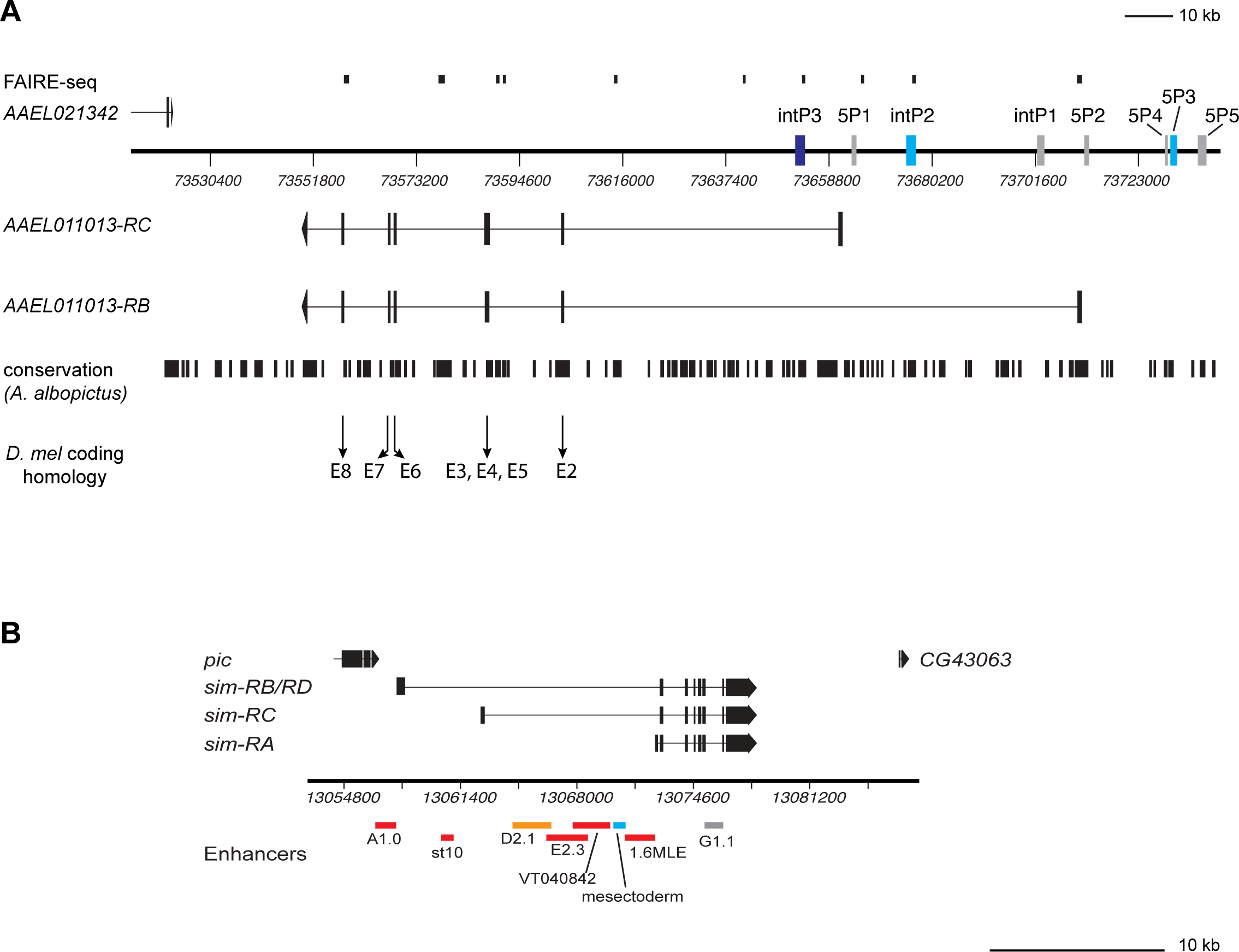
The *A. aegypti* and *D. melanogaster sim* loci. Note that the two maps are at different scales. (A) The *A. aegypti sim* locus (Vectorbase gene AAEL011013). The two annotated transcripts are shown along with positions of the sequences tested for enhancer activity in this paper. Sequences with no embryonic reporter gene activity are shown in gray, midline activity in cyan, and ectopic activity in dark blue. Positions of FAIRE peaks (from (46)), conservation with *A. albopictus*, and exon conservation with *D. melanogaster* are shown. Conservation with *A. albopictus* was assessed by Blast2Seq run on the NCBI Blast server (70) using word size = 11, match/mismatch = 2,-3 and gapcosts = 5,2. (B) The *D. melanogaster sim* locus showing the annotated transcripts and a subset of the known *sim* enhancers. Enhancers with midline activity are in red, early (mesectoderm stage) activity in cyan, and weak midline activity in orange. Sequence *G1.1*, shown in gray, has conflicting reports in the literature about its midline activity but in our hands lacks midline expression.

**Supplemental Figure S4:**
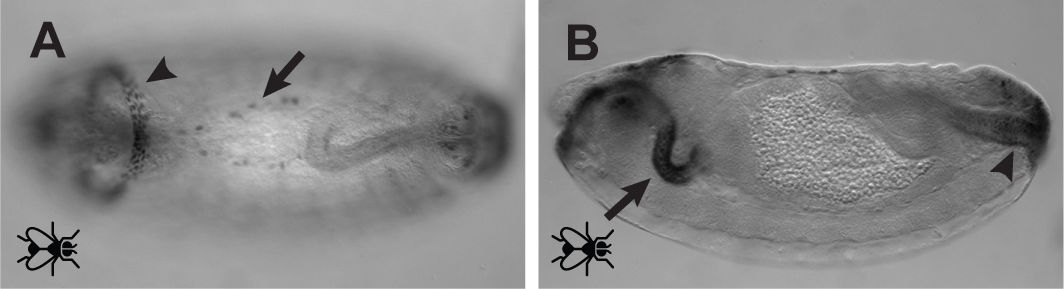
*intP3* reporter gene expression. All observed *intP3* activity is ectopic with respect to *sim*. (A) Dorsal view of a stage 14 embryo showing reporter gene expression in the developing dorsal vessel (arrowhead) and in a ring of cells in the anterior region of the embryo, provisionally identified as atrium precursors. (B) Sagittal view of a stage 15 embryo showing prominent expression in the foregut (arrow) as well as weaker expression in the hindgut (arrowhead).

**Supplemental Figure S5:**
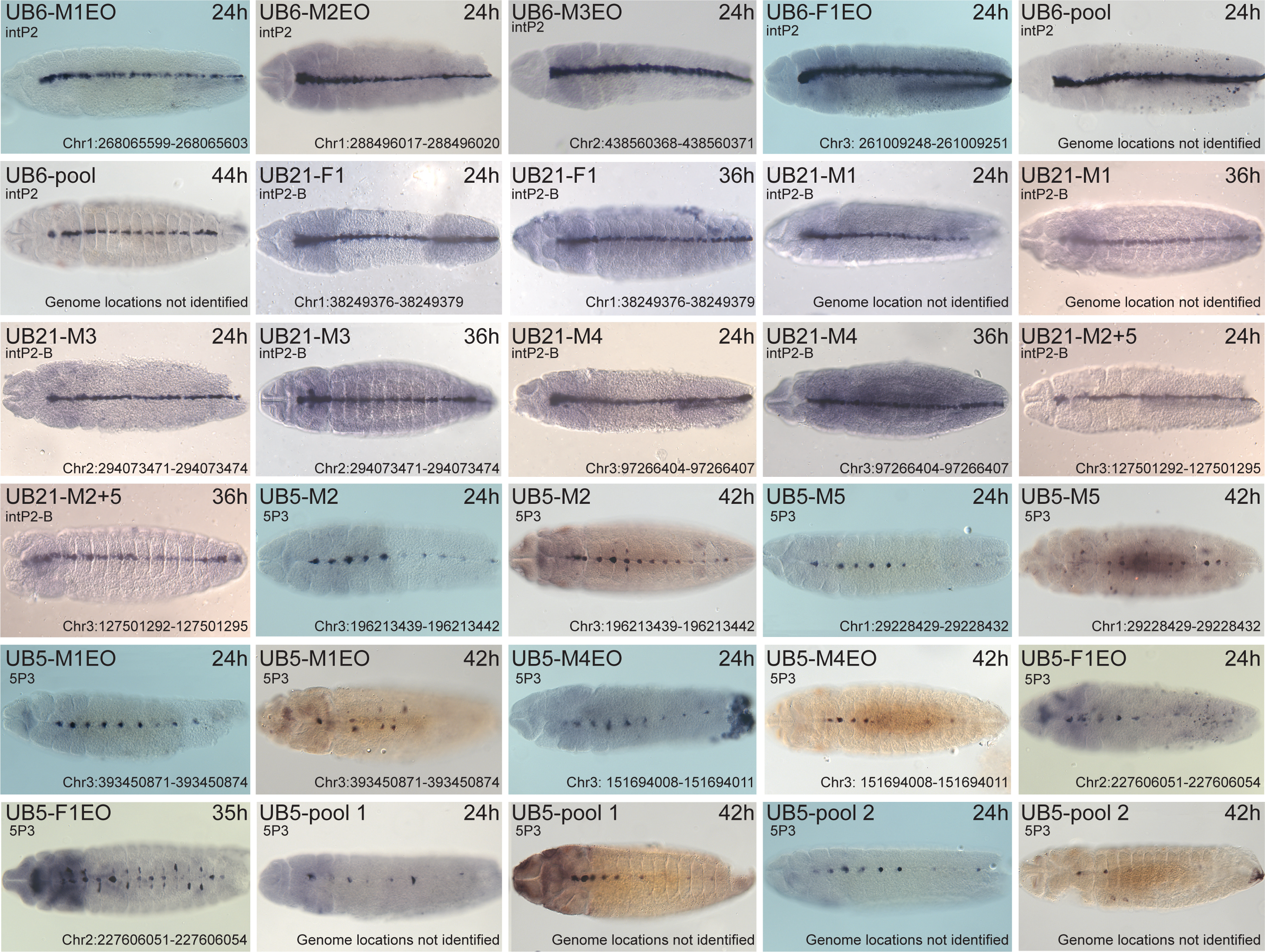
Additional transgenic lines. Immunohistochemical staining of the eGFP reporter in additional lines of transgenic *A. aegypti* for the *intP2, intP2B,* and *5P3* CRMs. No qualitative differences in expression pattern were observed among the multiple independent lines for each reporter construct. All embryos are oriented ventral side up with anterior to the left.

**Supplemental Figure S6:**
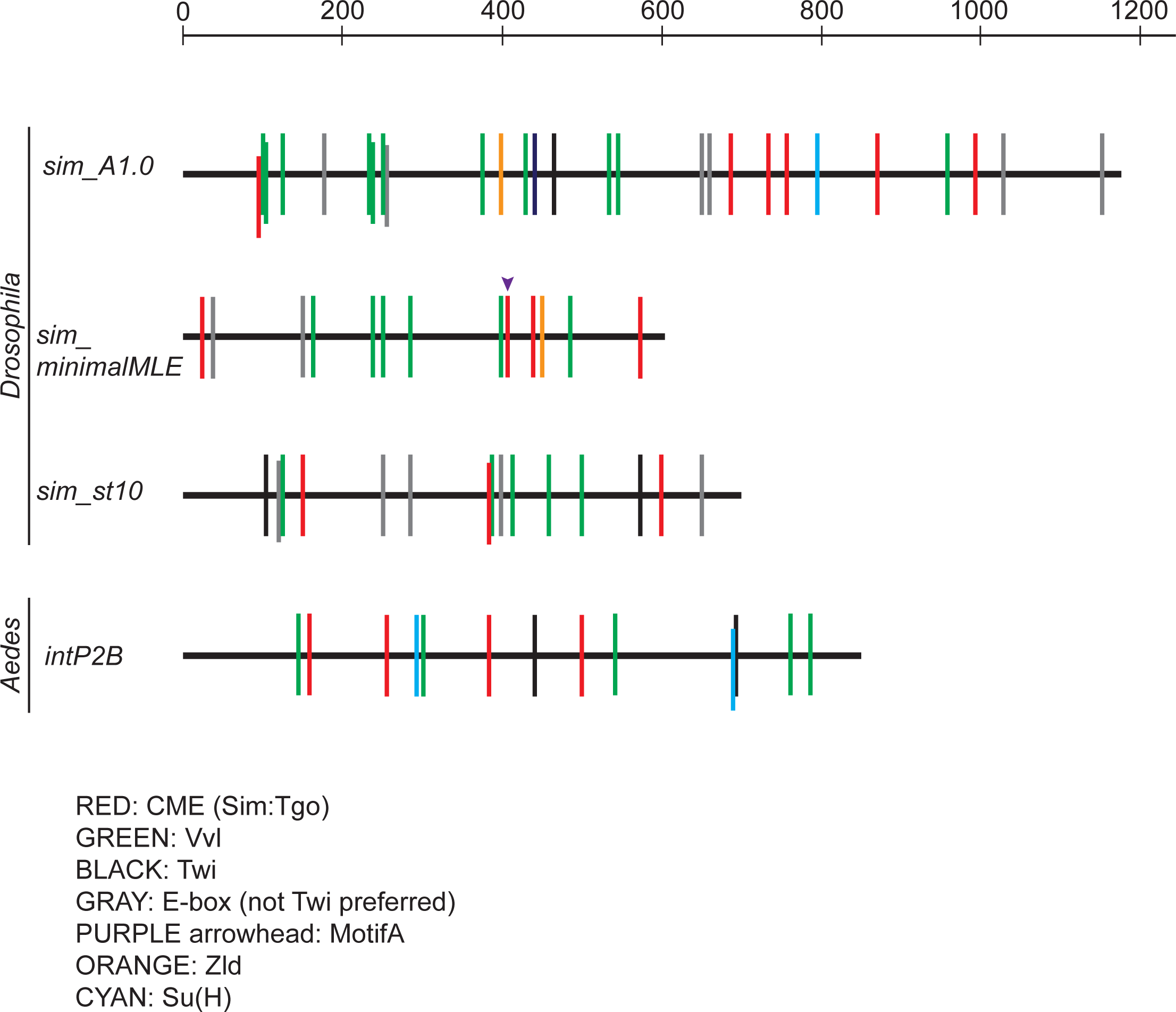
Known *D. melanogaster* and *A. aegypti sim* enhancers. Sequences of known *D. melanogaster sim* enhancers were obtained from REDfly (71) and analyzed for a subset of putative transcription factor binding sites using consensus sequences as described in Pearson et al. (25). *sim_minimalMLE* is a subfragement of *sim_1.6MLE* (Fig. S3B). The 2.3 kb *sim_E2.3* sequence is not included. *A. aegypti intP2B* is from the current study. The scale along the top indicates size in basepairs.

**Supplemental Figure S7:**
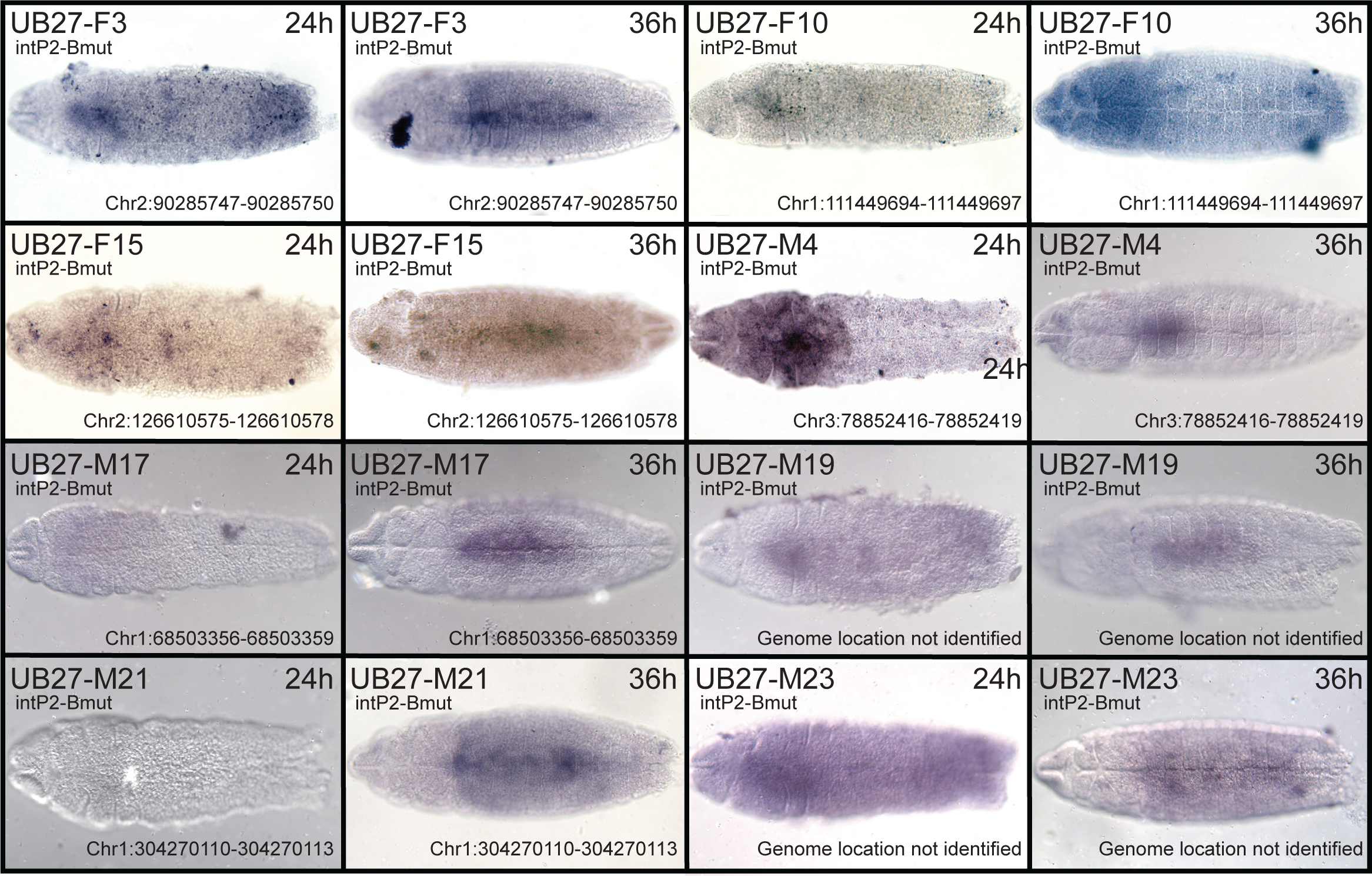
Additional transgenic lines with the *intP2Bmut* mutated enhancer in *A. aegypti*. Immunohistochemical staining of the eGFP reporter in seven additional lines of transgenic *A. aegypti* for the *intP2Bmut* CRM (line UB27-F15 is pictured in Fig. 2L, M). No qualitative differences in expression pattern were observed among the multiple independent lines. All embryos are oriented ventral side up with anterior to the left.

**Supplemental Figure S8:**
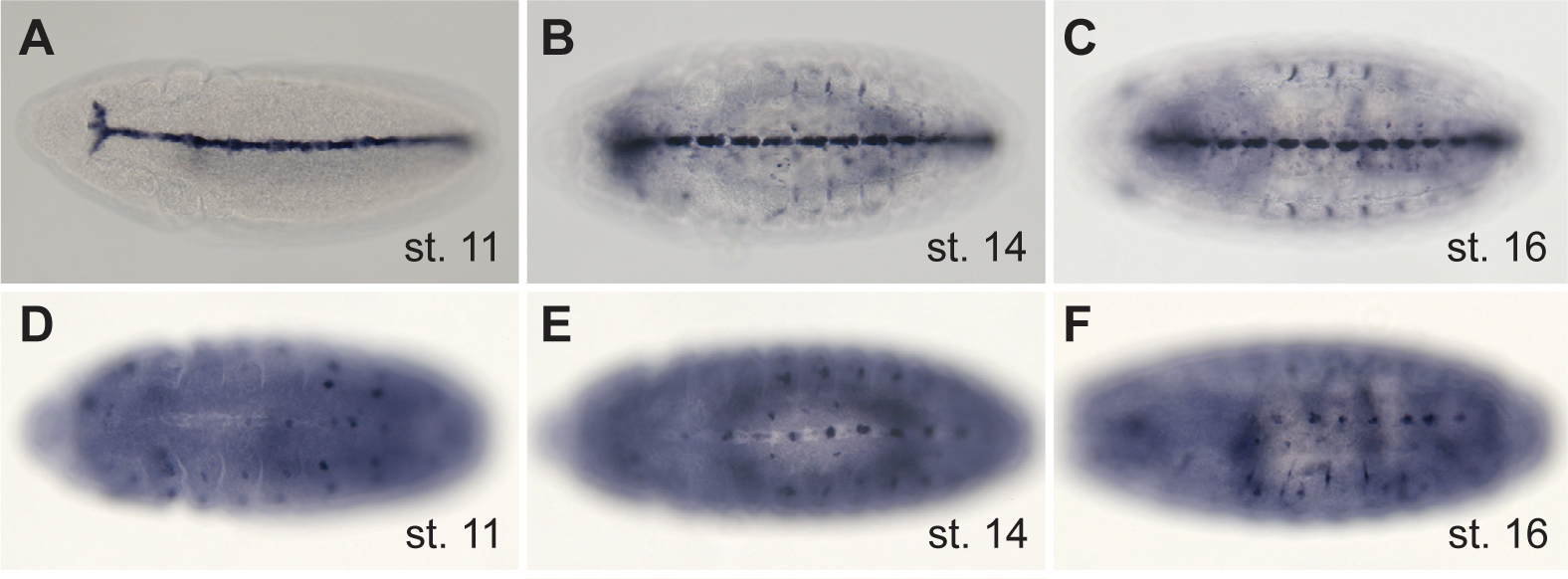
*A. albopictus* enhancer activity in transgenic *Drosophila*. (A-C) The *A. albopictus intP2B* CRM is active in the expected pattern in *Drosophila* embryos at stages 11, 14, and 16, respectively. (D-F) The *A. albopictus 5P3* CRM is active in the expected pattern in *Drosophila* embryos at stages 11, 14, and 16, respectively. Some segments are out of focus. All embryos are oriented ventral side up with anterior to the left.

**Supplemental Figure S9:**
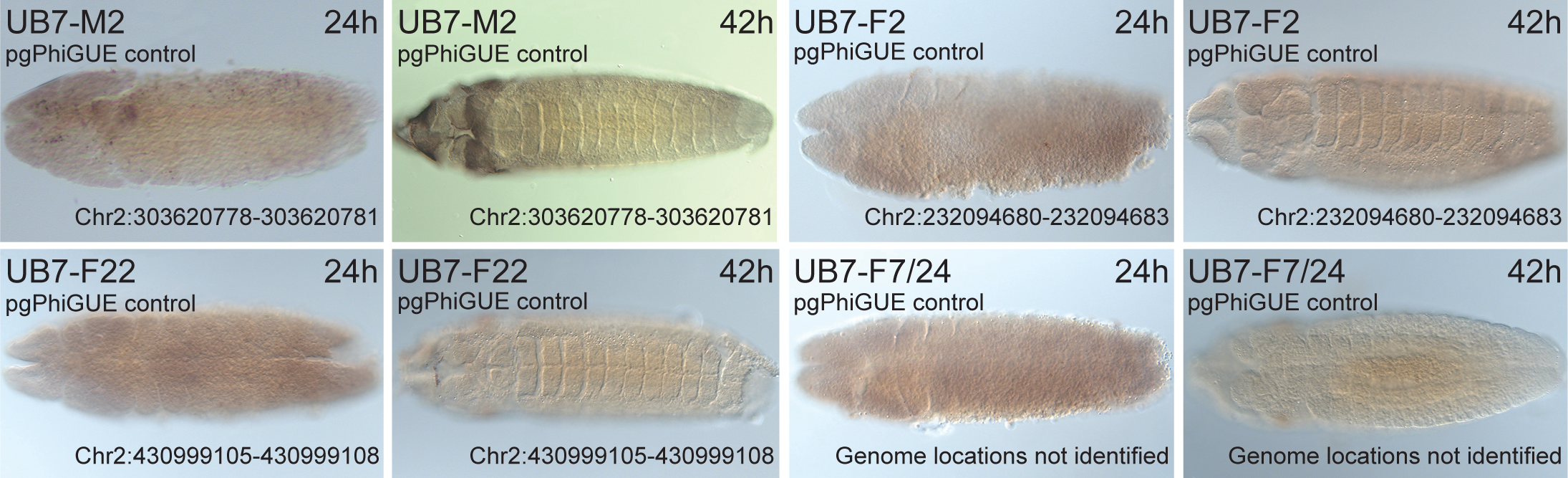
Vector control lines. Immunohistochemical staining of the eGFP reporter for a negative control reporter (*pgPhiGUE*_LANDR, courtesy of Kevin Deem) demonstrating the absence of vector-dependent reporter gene activity. All embryos are oriented ventral side up with anterior to the left.

**Supplemental Figure S10:**
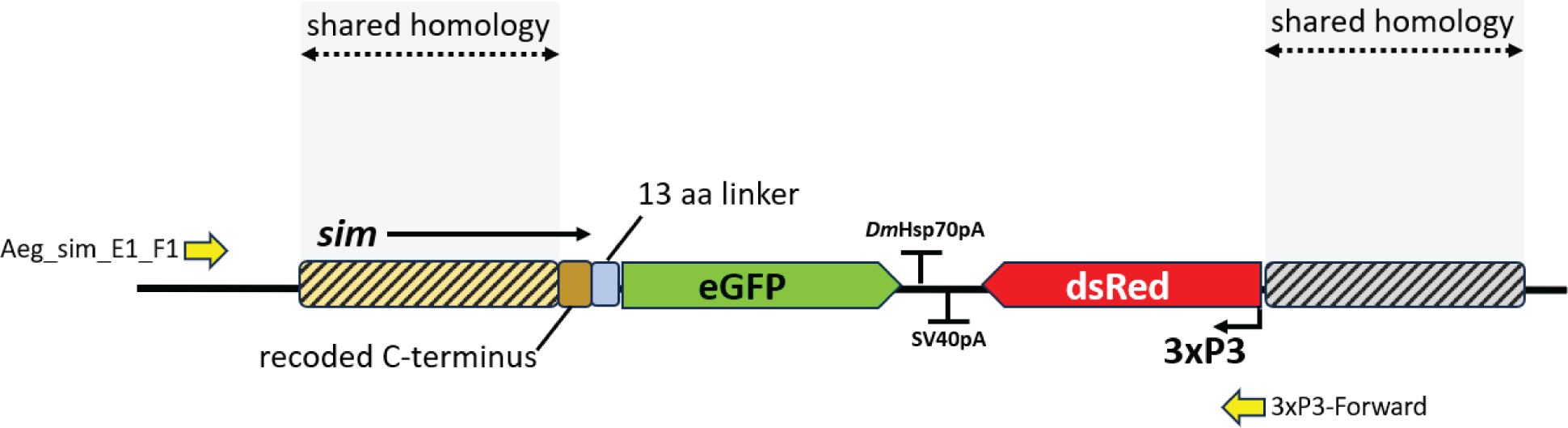
Schematic showing the integration of the sim::eGFP fusion construct along with the 3xP3-dsRed eye marker. Sequencing through the region with primers *Aeg_sim_E1_F1* and *3xP3-Forward* confirmed the correct integration.

