## Supplemental Files for "Conserved and novel enhancers in the *Aedes aegypti single-minded* locus recapitulate embryonic ventral midline gene expression"

**Supplemental Figure S10: Schematic showing the integration of the sim::eGFP fusion construct along with the 3xP3-dsRed eye marker.** Sequencing through the region with primers *Aeg_sim_E1_F1* and *3xP3-Forward* confirmed the correct integration.

**Table S1:** Primers for transgenic fly and mosquito lines

| Fragment | Vector[a] | Forward primer[b] | Reverse Primer[c] |
|---|---|---|---|
| *intP1* | pLacZattB | tcgcaggcctcgcaccggtg*aattc*ACTATGTCGATAACTGGTAC | ccggcgctctagaggtaccc*tcgag*CTTAAGGTCAAATACAAATGG |
| *intP2* | pLacZattB & pgPhiGUE | tcgcaggcctcgcaccggtg*aattc*CAATTTAGTAGTTTTTTTTTGCAATTTC | ccggcgctctagaggtaccc*tcgag*GTGGTAAAAATCAATAGATTCC |
| *intP2A* | pLacZattB | tcgcaggcctcgcaccggtg*aattc*CCGTGCGTCCGTTAACAAATC | ccggcgctctagaggtaccc*tcgag*CGTATCTGGACCACTGTG |
| *intP2B* | pLacZattB | tcgcaggcctcgcaccggt*gaattc*GCTGCTAAAATGCCGAGTC | ccggcgctctagaggtaccc*ctcgag*TATTACCCCCTCCCCTCG |
| *intP3* | pLacZattB | tcgcaggcctcgcaccggtg*aattc*TGAAAATATTTGGAAACGACTTAC | ccggcgctctagaggtaccc*tcgag*TCTCAATTATACGCGGCG |
| *5P1* | pLacZattB | tcgcaggcctcgcaccggtg*aattc*ATATGCAAATCATATTCTGATAAAATTTTATTATATTT | ccggcgctctagaggtaccc*tcgag*ACCAGAATGATAATTAAAATATAAATGTC |
| *5P2* | pLacZattB | tcgcaggcctcgcaccggtg*aattc*CTCATCATGAAATTTCCAGATAG | ccggcgctctagaggtaccc*tcgag*TATTCTACTAACACCAAACTAG |
| *5P3* | pLacZattB & pgPhiGUE | tcgcaggcctcgcaccggtg*aattc*ATTAAATAAACTTTACCGAATGATC | ccggcgctctagaggtaccc*tcgag*AATAAAAATGGCGACCGATG |
| *5P3A* | pLacZattB | tcgcaggcctcgcaccggtg*aattc*GCGCCAAGCCAAATTTAC | ccggcgctctagaggtaccc*tcgag*CACCGTGCCAGAGAACATC |
| *5P3B* | pLacZattB | tcgcaggcctcgcaccggtg*aattc*AAGCGAAAGGGTGTTCAAG | ccggcgctctagaggtaccc*tcgag*GGCAATGGAGGAGCAATG |
| *5P3D* | pLacZattB | tcgcaggcctcgcaccggtg*aattc*TTCCATTCAACGTTTGCC | ccggcgctctagaggtaccc*tcgag*AGTCTATGGTAGCACTGC |
| *5P3E* | pLacZattB | tcgcaggcctcgcaccggtg*aattc*TGAGTTCTAAGAGCATGTTCC | ccggcgctctagaggtaccc*tcgag*TAAATCGCCAGGTGAGCC |
| *5P3F* | pLacZattB | tcgcaggcctcgcaccggtg*aattc*TTCAACGTTTGCCCGGAAAC | ccggcgctctagaggtaccc*tcgag*ATGGCGACCGATGGTTGTTAC |
| *5P3G* | pLacZattB | tcgcaggcctcgcaccggtg*aattc*TCCTTTCAAGTCCTAAAAGAAAAGAACATTGTTAC | ccggcgctctagaggtaccc*tcgag*CGGTTTCCGGGCAAACGTTG |
| *5P3H* | pLacZattB | tcgcaggcctcgcaccggtg*aattc*ACCCTTAAATAAATCCCCAATTTG | ccggcgctctagaggtaccc*tcgag*TCTTTTCTTTTAGGACTTGAAAG |
| *5P3I* | pLacZattB | tcgcaggcctcgcaccggtg*aattc*TGAGTTCTAAGAGCATGTTCC | ccggcgctctagaggtaccc*tcgag*GCAAATTGGGGATTTATTTAAG |
| *5P4* | pLacZattB | tcgcaggcctcgcaccggtg*aattc*TGACATTTGTAGCATTGTCAC | ccggcgctctagaggtaccc*tcgag*ATAGCAGGACCGTACCAG |
| *5P5* | pLacZattB | tcgcaggcctcgcaccggtg*aattc*ATGAAGCAAGCACCTCGG | ccggcgctctagaggtaccc*tcgag*CCGACCTTCCACAGCTATC |

[a] The *pLacZattB* vector was used to make transgenic *D. melanogaster*, while the *pgPhiGUE* vector was used to make transgenic *A. aegypti*

[b] Lowercase letters are specific to *pattBLacZ*, italics indicate the restriction site for *EcoRI*, and uppercase letters are specific to the amplified *A. aegypti* sequence

[c] Lowercase letters are specific to *pattBLacZ*, italicized indicate the restriction site for *XhoI*, and uppercase letters are specific to the amplified *A. aegypti* sequence

**Table S2.** Genomic insertion sites for the transgenic *Ae. aegypti* lines

| Construct | Line prefix | Line | Genomic insertion site (LVP genome version: AaegL5.3) |
|---|---|---|---|
| *5P3* | UB5 | M1EO | Chr3:393450871-393450874 |
| | | M5 | Chr1:29228429-29228432 |
| | | F1EO | Chr2:227606051-227606054 |
| | | M2 | Chr3:196213439-196213442 |
| | | M4EO | Chr3:151694008-151694011 |
| | | pool #1 | pooled line* |
| | | pool #2 | pooled line* |
| *intP2* | UB6 | M1EO | Chr1:268065599-268065603 |
| | | M2EO | Chr1:288496017-288496020 |
| | | M3EO | Chr2:438560368-438560371 |
| | | F1EO | Chr3:261009248-261009251 |
| | | pool #1 | pooled line* |
| *pgPhiGUE_LANDR* | UB7 | F2 | Chr2:232094680-232094683 |
| | | M2 | Chr2:303620778-303620781 |
| | | F22 | Chr2:430999105-430999108 |
| | | F7, F24 | pooled line* |
| *intP2B* | UB21 | F1 | Chr1:38249376-38249379 |
| | | M1 | Insertion not identified |
| | | M2+5 | Chr3:127501292-127501295 |
| | | M3 | Chr2:294073471-294073474 |
| | | M4 | Chr3:97266404-97266407 |
| *intP2Bmut* | UB27 | F3 | Chr2:90285747-90285750 |
| | | F10 | Chr1:111449694-111449697 |
| | | F15 | Chr2:126610575-126610578 |
| | | M4 | Chr3:78852416-78852419 |
| | | M17 | Chr1:68503356-68503359 |
| | | M19 | Insertion not identified |
| | | M21 | Chr1:304270110-304270113 |
| | | M23 | Insertion not identified |

*Genomic insertion sites were not determined for pooled samples with multiple piggyBac insertion sites.

**Table S3.** Primers and gBlocks used to generate the *sim*::eGFP fusion line of *Ae. aegypti.*

| Primer/gBlock | Sequence (5’ -> 3’) |
|---|---|
| *sim-eGFP-don-MT* | caattgggcccGATCCaacaccagcagccggggtacgatgagataaatCAAcacttcggtctgaactgtttcaacaata<br>ataataccagcaacggcaaTggcgccattcctgccaacggtcatttgaatggtggtctcaacagtcacactctacatcaccag<br>cacagtgccaacagcaataacggaagcaacagtagcagcgccagtgctgccaacagtaataacaacaatcacgacaatggc<br>ttcatggatgagttcaaagtaagccaacagcatcaccatcatccccagcagcagcaacagcatgtcgtccctcagcagcacca<br>tcatccgcaccaccaacaacatcaccaccacagtcacaacggggcagggggaggacctcagtacaccagcgtgattgtgga<br>gccgCAAaattaccacctaccagtaactcagaatgagttcgttcacggatcggctggatcggctgctggatcgggagaattc<br>caattctcgagctatcatctgccggtcacgcaaaacgaatttgtgcattaggcgccgccattttccccctcctttccctccgggaa<br>ggagtttatgcatcgtcagtgggcaaaaactgactgaaagcagcaaagaattAagcagtttgtgctgccgttgattgtgaataa<br>cgagttagactatttttgaaaaggaaaacctctaaagggaatcgattgatactagtttcaaccattgcgagttgatctgaggggct<br>catttagtcccaaattcccttgcattttagtagtttttctagaacttattatagttttgcgtcttgtttcatgtgattattgtttcgaaatgtttt<br>catattgtacagagatacagtgctggacattattatgtaatGtcgtttttcacacaaaatccacatctccatacattcaacgcactgt<br>gcggcTGTGCTGTGCGACTCCGTCGAGTCGACCAACATAGTTGAAACAAATTG<br>AATATTTAATTGATCGTTATAGGAATGGTGTTAGATGAGTCATCCTTTACAGTA<br>AGCACATACAGTATTATAATTGAAGATCGTCGGCAGATAGGTGTGTAGGGTAG<br>AGTATCAGCAATAAGTTGGGACGTTTGACTTTTTGTAGGTAGACAAAAACTA<br>AACtttttttCGCTTCTCTATGTGTGCCCCTGGGTAGCGTTCCGTTCCGATTGGGGT<br>GCGAACGAATGAAATCGCCCATCGAGTTGATACGTCCATCCATCGCTAGAACC<br>GCGTTCGCTGTAaAAGACTATATAAGAGCAGAGGCAAGAGTAGTGAAATaccgg<br>cagatgatagttttgGTTTCAGAGCTATGCTGGAAACAGCATAGCAAGTTGAAATAAG<br>GCTAGTCCGTTATCAACTTGAAAAAGTGGCACCGAGTCGGTGCTTTTTTTTTg<br>ctagc |
| *eGFP-fus-EcoRI-F* | caattGAATTCatggtgagcaagggcgag |
| *3xP3-fus-XhoI-R* | caattCTCGAGtgatcgcacggttcccac |
| *Aeg_sim_E7_F1* | CCAGCAAACAAGAAGCGCACG |
| *3xP3-Forward* | aattcgagctcgcccgg |

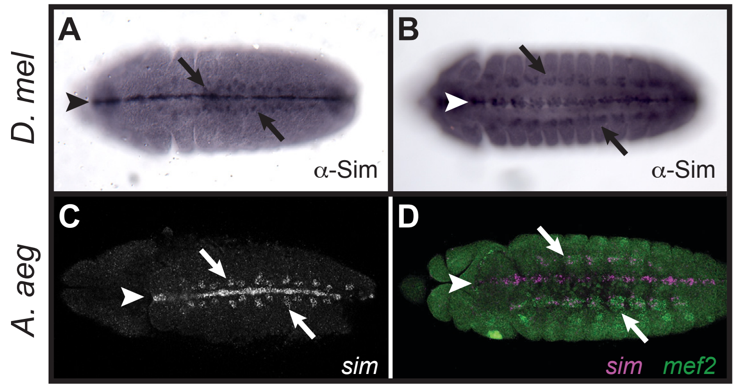

D. mel
A. aeg
A
B
C
D
α-Sim
α-Sim
sim
sim mef2

### Supplemental Figure 2

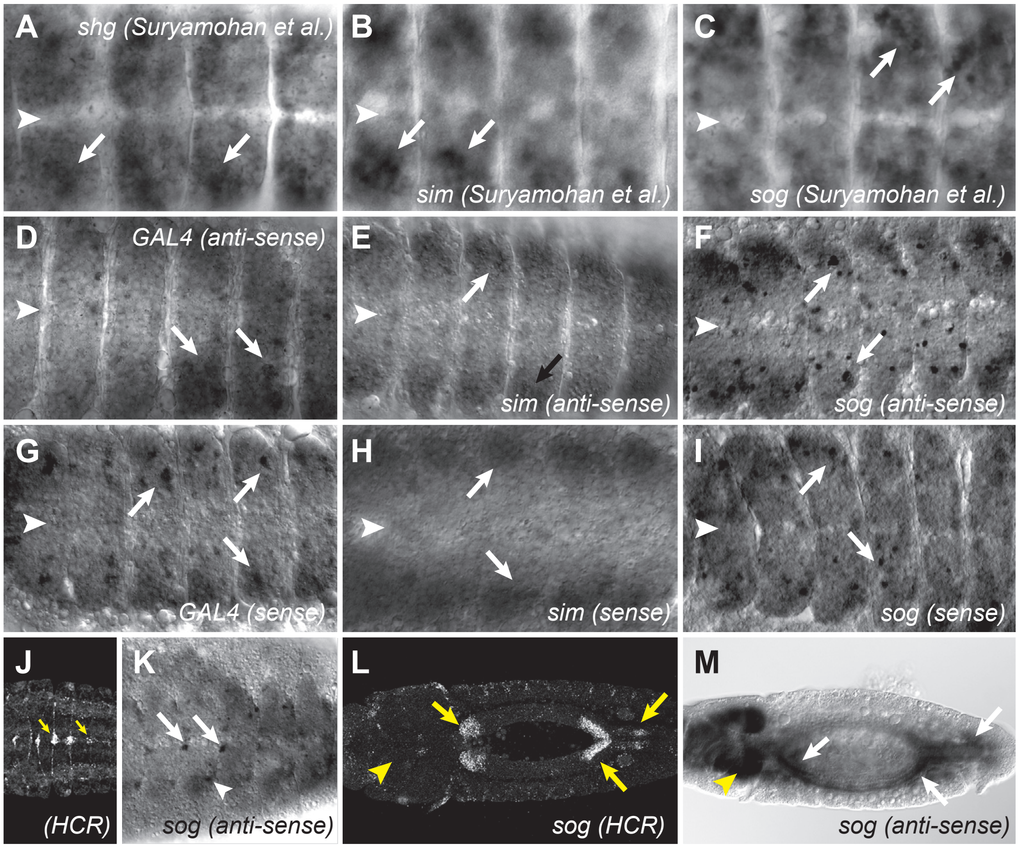

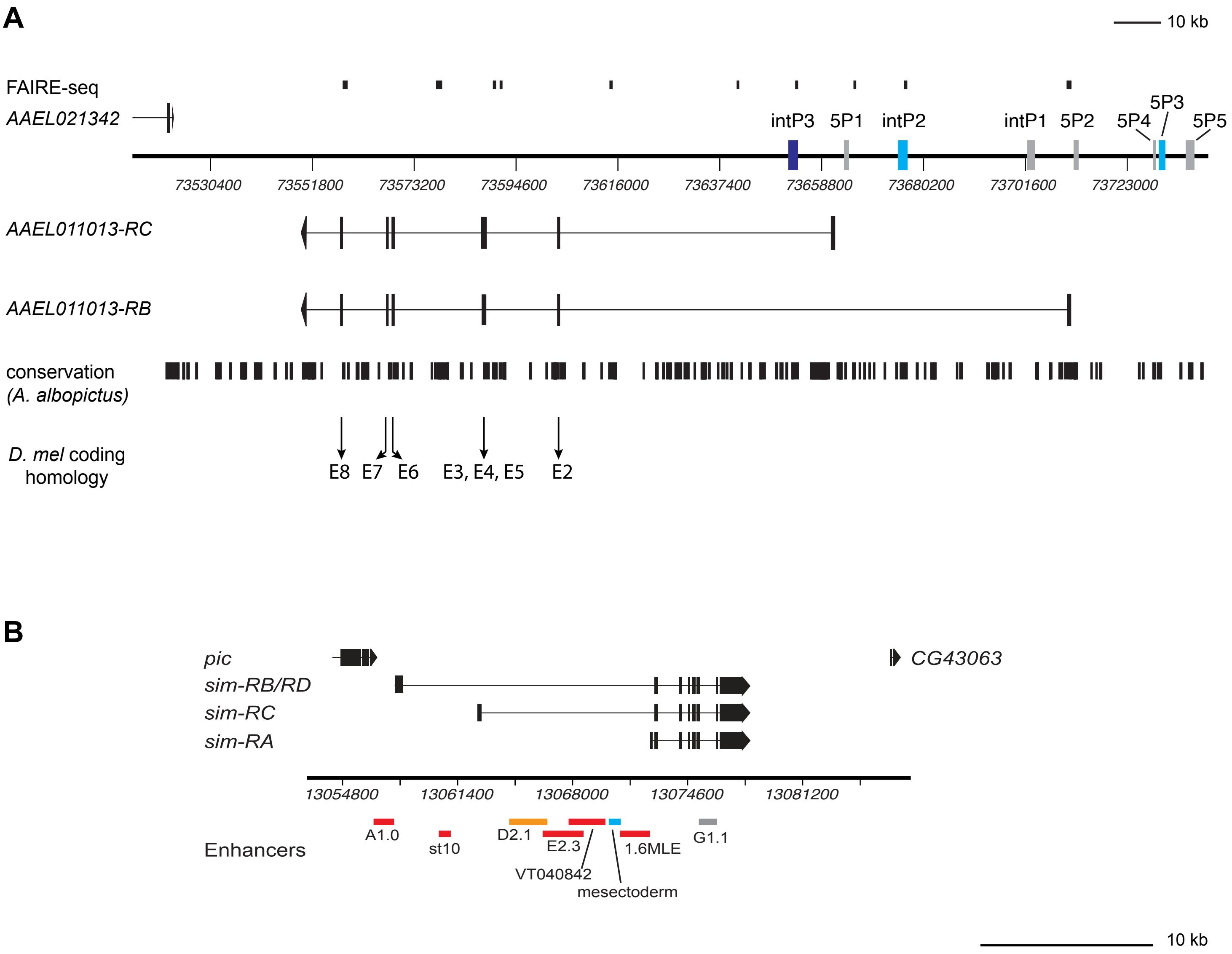
A
10 kb
FAIRE-seq
AAEL021342
intP3
5P1
intP2
intP1
5P2
5P4
5P3
5P5
73530400
73551800
73573200
73594600
73616000
73637400
73658800
73680200
73701600
73723000
AAEL011013-RC
AAEL011013-RB
conservation
(A. albopictus)
D. mel coding homology
E8
E7
E6
E3, E4, E5
E2
B
pic
CG43063
sim-RB/RD
sim-RC
sim-RA
13054800
13061400
13068000
13074600
13081200
Enhancers
A1.0
st10
D2.1
E2.3
VT040842
mesectoderm
1.6MLE
G1.1
10 kb

Supplementary Figure 4

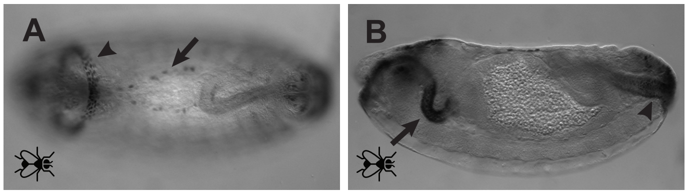

Schember et al. 2024. Supplemental Figure S5

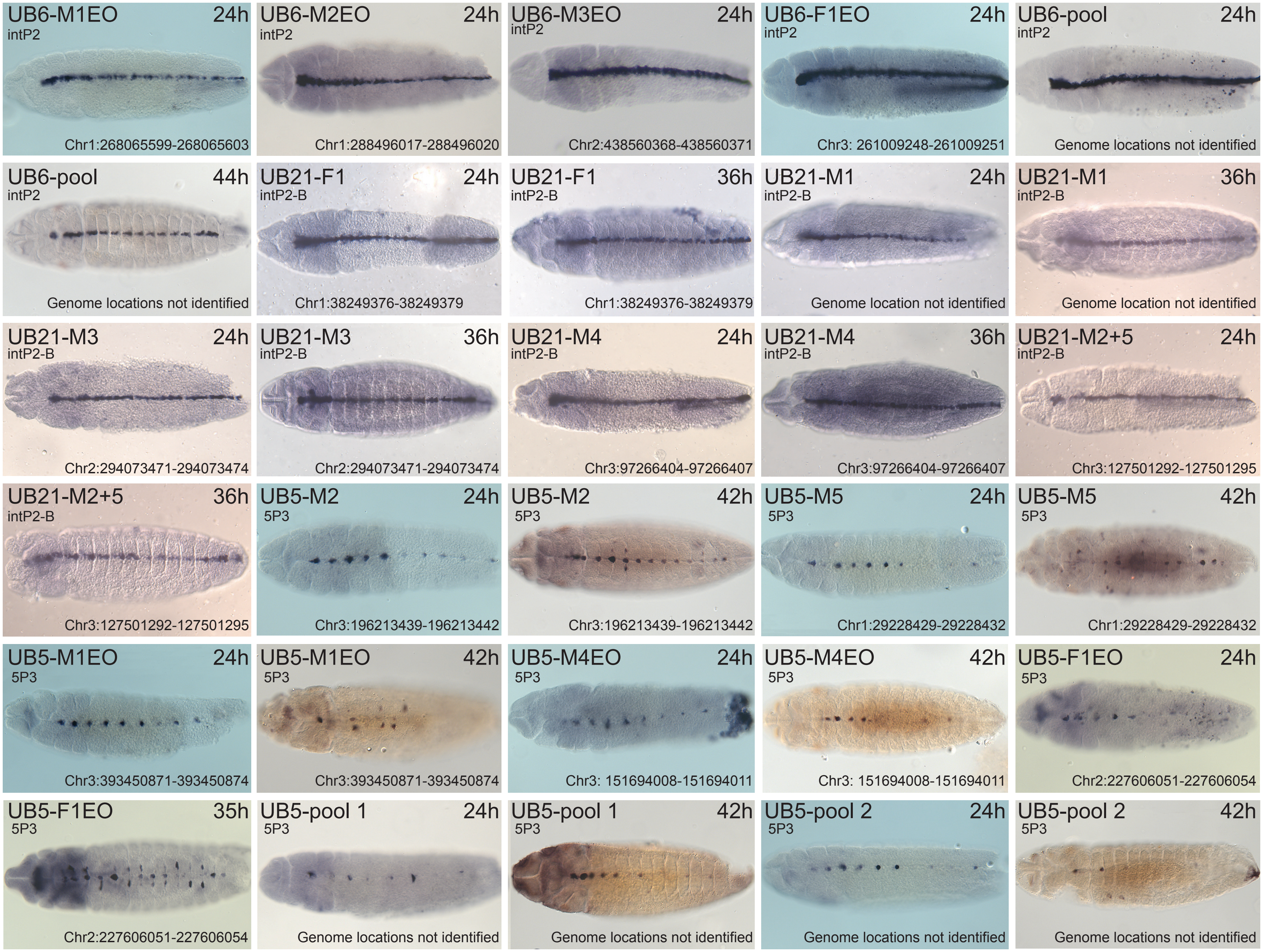

Supplemental Figure 6

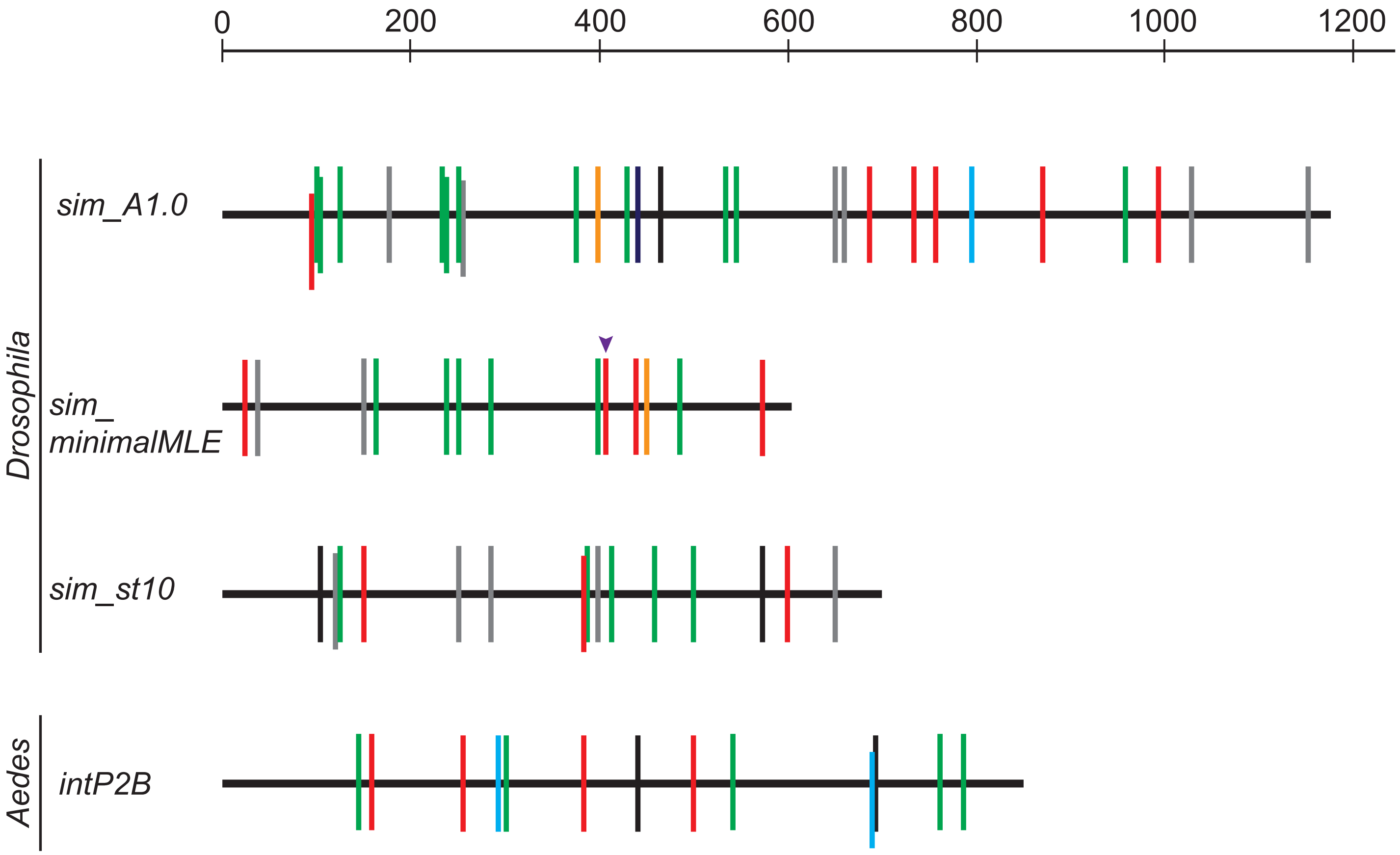


RED: CME (Sim:Tgo)
GREEN: Vvl
BLACK: Twi
GRAY: E-box (not Twi preferred)
PURPLE arrowhead: MotifA
ORANGE: Zld
CYAN: Su(H)

Schember et al. 2024 Supplemental Figure S7

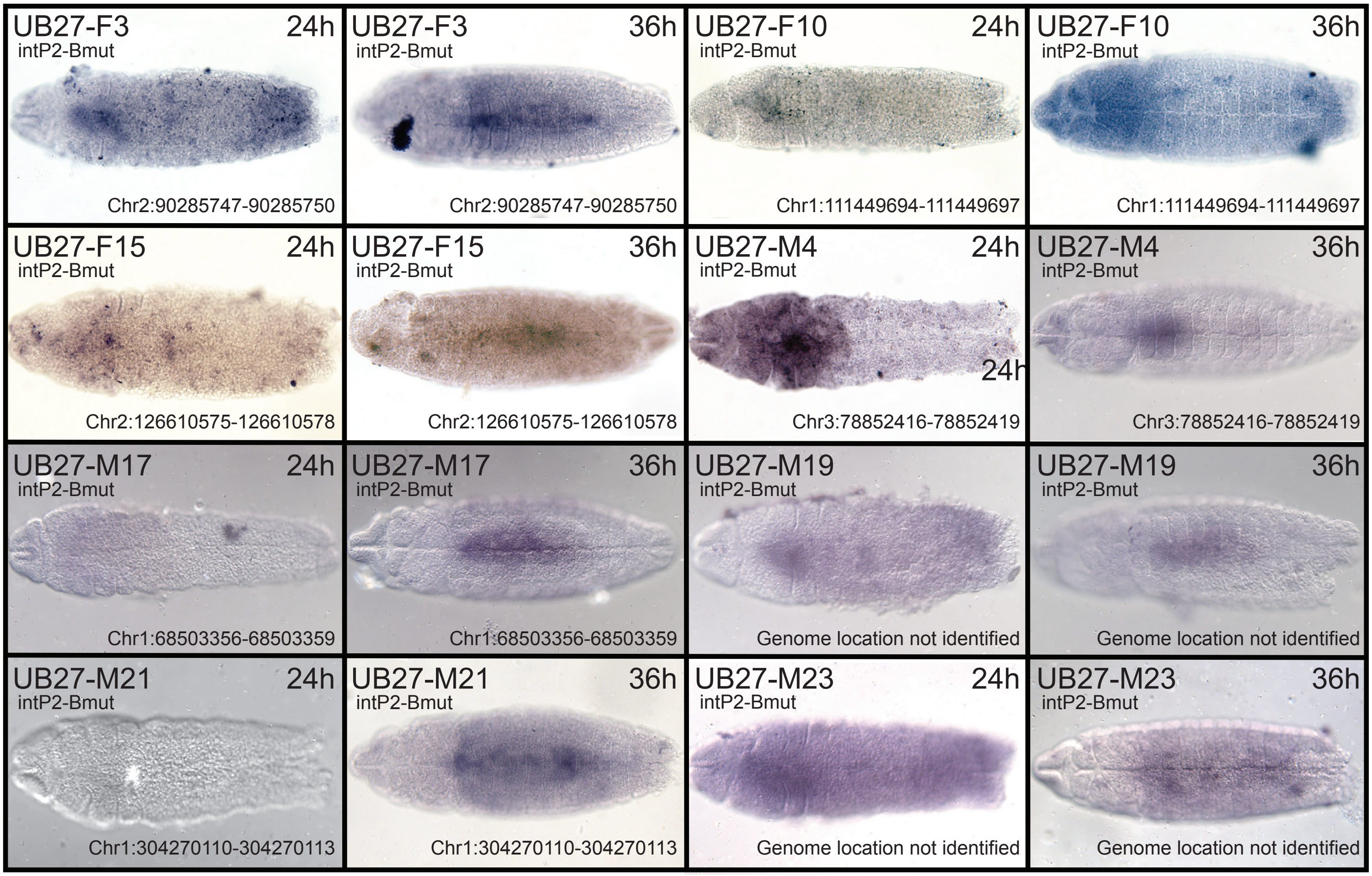

Schember et al. 2024. Supplemental Figure S8

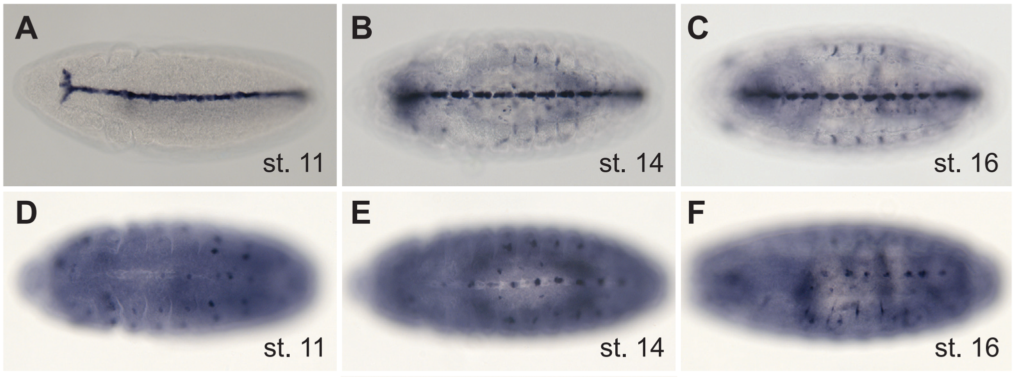

Schember et al. Supplemental Figure S9

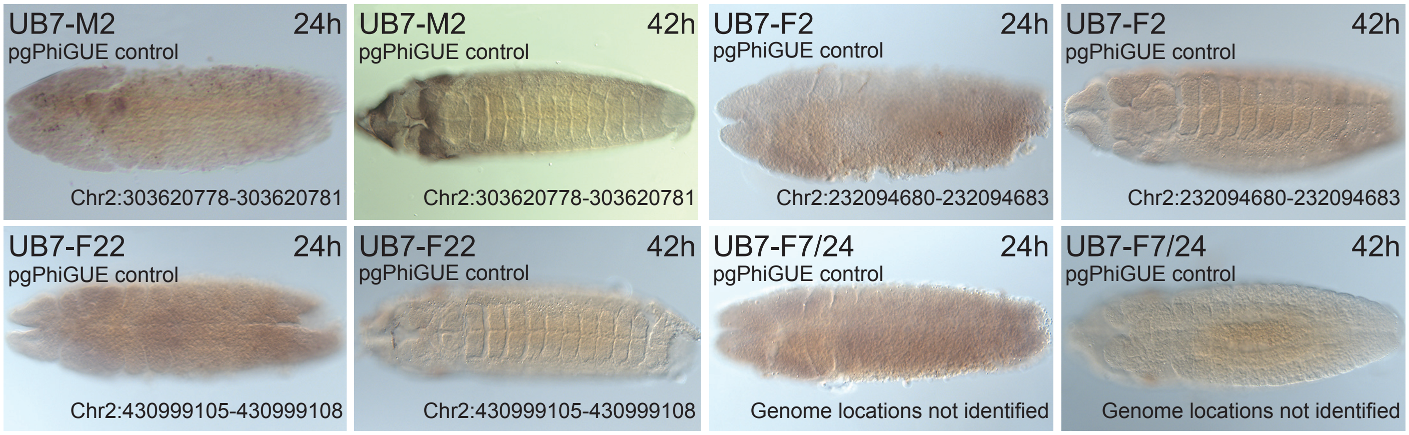

Figure S10

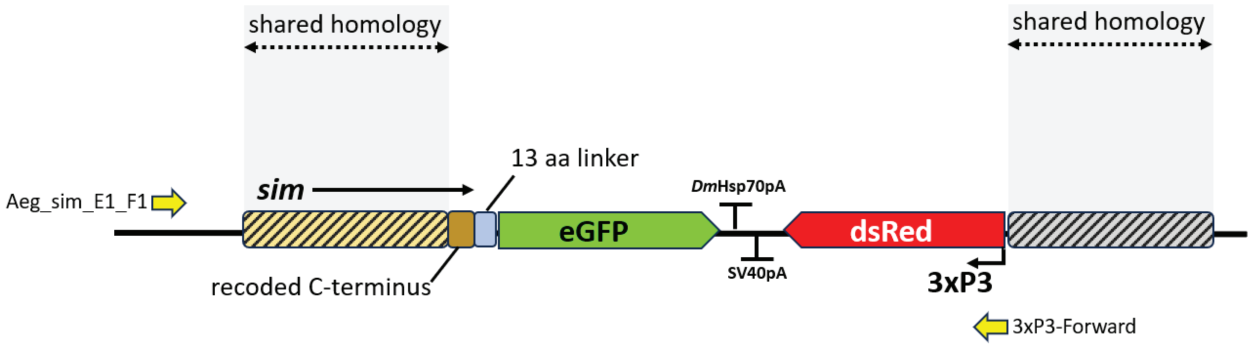
shared homology
shared homology
13 aa linker
Aeg_sim_E1_F1
sim
DmHsp70pA
eGFP
dsRed
SV40pA
recoded C-terminus
3xP3
3xP3-Forward
